# Comprehensive mapping of cell fates in microsatellite unstable cancer cells support dual targeting of WRN and ATR

**DOI:** 10.1101/2023.07.28.550976

**Authors:** Dali Zong, Natasha C. Koussa, James A. Cornwell, Ajith V. Pankajam, Michael J. Kruhlak, Nancy Wong, Raj Chari, Steven D. Cappell, André Nussenzweig

## Abstract

Addiction to the WRN helicase is a unique vulnerability of human cancers with high levels of microsatellite instability (MSI-H). However, while prolonged loss of WRN ultimately leads to cell death, little is known about how MSI-H cancers initially respond to acute loss of WRN, knowledge that would be helpful for informing clinical development of WRN-targeting therapy, predicting possible resistance mechanisms, and identifying useful biomarkers of successful WRN inhibition. Here, we report the construction of an inducible ligand-mediated degradation system wherein the stability of endogenous WRN protein can be rapidly and specifically tuned, enabling us to track the complete sequence of cellular events elicited by acute loss of WRN function. We find that WRN degradation leads to immediate accrual of DNA damage in a replication-dependent manner that curiously did not robustly engage checkpoint mechanisms to halt DNA synthesis. As a result, WRN-degraded MSI-H cancer cells accumulate DNA damage across multiple replicative cycles and undergo successive rounds of increasingly aberrant mitoses, ultimately triggering cell death. Of potential therapeutic importance, we find no evidence of any generalized mechanism by which MSI-H cancers could adapt to near-complete loss of WRN. However, under conditions of partial WRN degradation, addition of low dose ATR inhibitor significantly increased their combined efficacy to levels approaching full inactivation of WRN. Overall, our results provided the first comprehensive view of molecular events linking upstream inhibition of WRN to subsequent cell death and suggested a potential therapeutical rationale for dual targeting of WRN and ATR.

## Introduction

Microsatellite instability (MSI) is a pathological condition of pervasive hypermutation that occurs at short tandem repeat DNA sequences within the genome, caused by defects in cellular mismatch repair (MMR) (Olave & Graham, 2022). MSI has been observed in many types of human cancers, most commonly in colorectal, endometrial, and gastric adenocarcinomas. Tumors with high levels of microsatellite instability (MSI-H) tend to respond well to immunotherapy (Taieb *et al*, 2022) (Jin & Sinicrope, 2022) but may be relatively less responsive to conventional chemotherapy. Nevertheless, chemotherapy remains the standard of care for patients with MSI-H cancers (Taieb *et al*., 2022).

In the past four years, multiple studies have identified a unique addiction of human MSI-H cancers to the RECQ family helicase WRN (Morales-Juarez & Jackson, 2022) (Picco *et al*, 2021) (van Wietmarschen *et al*, 2020) (Chan *et al*, 2019) (Behan *et al*, 2019) (Lieb *et al*, 2019) (Kategaya *et al*, 2019), uncovering one of the most striking examples of synthetic lethality since PARP inhibition was shown to specifically kill homologous recombination (HR) defective breast/ovarian cancers two decades ago (Farmer *et al*, 2005) (Bryant *et al*, 2005). This has led to a huge push from the pharmaceutical industry to develop clinical grade WRN inhibitors. Yet, despite the obvious benefit of targeting WRN, our understanding of how upstream WRN deficiency triggers downstream death of MSI-H cancers cells remains incomplete. We have shown earlier that MSI-H colorectal cancer cells contain unstable structure-forming TA repeats in their genome and that in the absence of WRN, these become fragile and ultimately break (van Wietmarschen Nathan & Nussenzweig, 2021) (van Wietmarschen *et al*., 2020). Our study, along with many others, thoroughly documented the long-term effects of WRN deficiency in MSI cancers using a combination of siRNA, shRNA and/or CRISPR-based approaches. However, due to the relatively slow and asynchronous nature of these reagents, so far it has not been possible to study the short-term, early effects of acute WRN loss in a manner that would mimic pharmacological inhibition.

To fill this gap in our knowledge and achieve a more complete elucidation of the cellular responses to WRN deficiency, we devised a strategy to engineer a degradable WRN protein encoded at its endogenous locus based on PROTAC technology. The modified WRN protein is functional and can be rapidly induced to degrade upon the addition of a small molecule ligand, dTAG-13 (Nabet *et al*, 2018). Using this model, we systemically mapped cell fates upon acute WRN loss. We show that the response of MSI cancer cells to WRN deficiency is biphasic and highly heterogeneous. In the acute phase, cells begin accruing replication-associated DNA damage almost as soon as WRN is degraded. Nevertheless, the majority of MSI cancer cells traverse two or more cell cycles before they die or become stably arrested (the chronic phase). Mechanistically, WRN degradation elicited a non-canonical DNA damage response (DDR) in MSI cancer cells, wherein repair factors such as 53BP1 and RAD51 were mobilized normally but a robust checkpoint response failed to initially engage, resulting in trans-cell cycle propagation and amplification of DNA damage. Of therapeutic relevance, we found that MSI cancers do not readily adapt to loss of WRN, which contrasts with the relative ease by which HR defective cancers become resistant to PARP inhibitors. Finally, we found that ATR inhibition mimicked WRN inactivation in MSI cancers and that low doses of ATR inhibitor can compensate for incomplete WRN degradation. As ATR inhibitors are already approved clinically, our work thus supports the use of ATR inhibitors either alone (at higher doses) or in tandem (at lower doses) with WRN inhibition to combat human MSI cancers.

## Results

### An endogenous, rapidly inducible system for controlled degradation of WRN

Prolonged loss of WRN is extremely toxic to human MSI-H cancer cells, but the sequence of proximal events triggered upon acute WRN loss, particularly signaling upstream of eventual cell death, remains largely unexplored. We therefore sought to engineer a new system capable of inducing rapid and controlled tuning of WRN protein levels, properties that would allow us to probe both short– and long-term consequences of WRN ablation.

To this end, we adapted PROTAC and CRISPR technologies to fuse an FKBP12^F36V^ degron domain to the N-terminus of WRN at its endogenous locus in three MSI-H cell lines and one MSS cell line (Fig 1A, B and Appendix Figure S1). We derived two types of clones: those that contained homozygous knock-in of FKBP-WRN and others that showed hemizygous knock– in with the second allele of WRN inactivated by a loss-of-function indel (deletion of ATG or nonsense mutation) (Appendix Figure S1B). In both cases, the FKBP-WRN protein was produced at a reduced level compared to unmodified WRN (Fig 1C). This level of WRN protein was nonetheless sufficient to promote survival of human MSI-H cancer lines.

**Fig 1.**
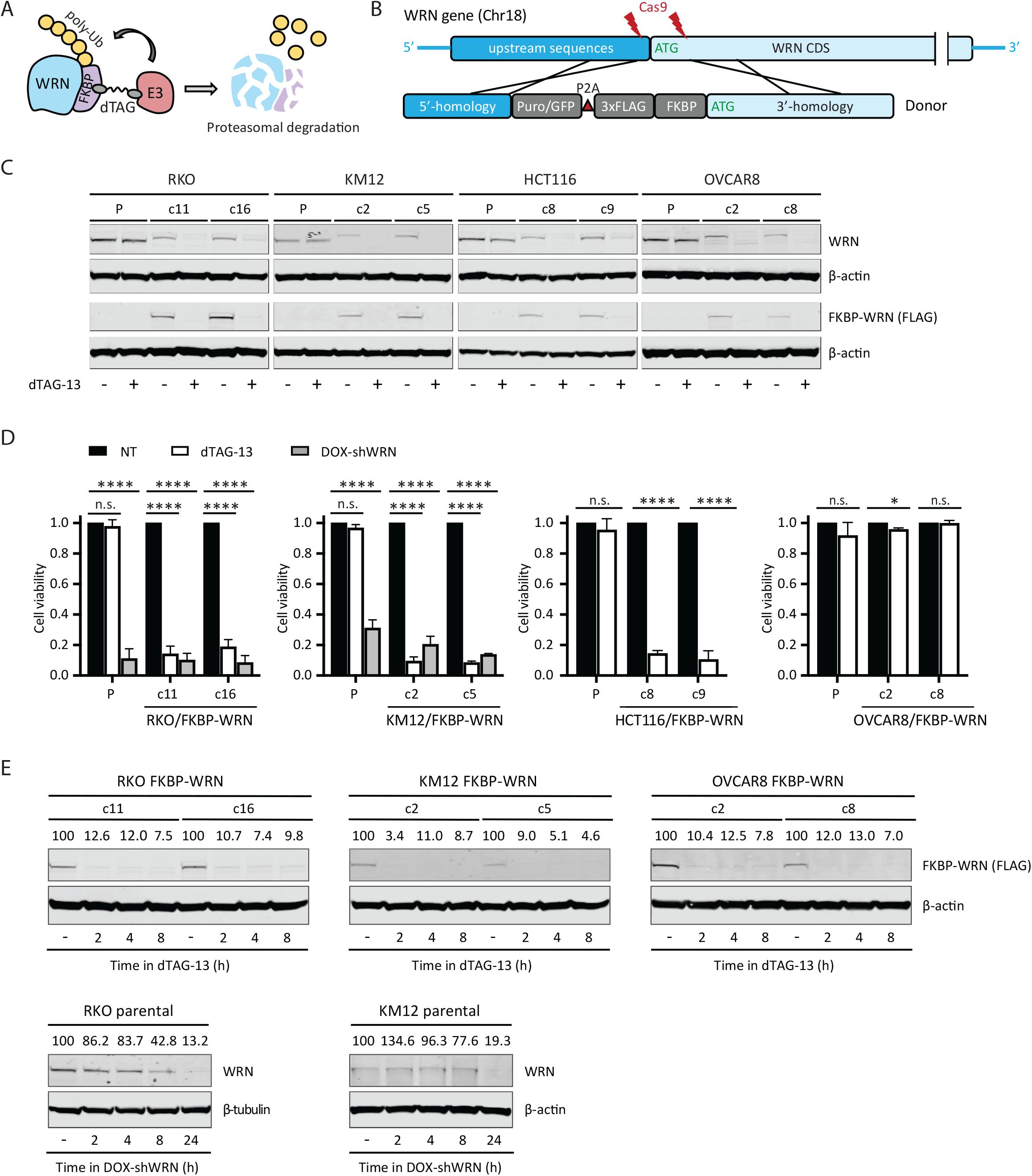
Generation of human cancer cells expressing degradable FKBP-WRN. (A) Schematics of the FKBP-WRN degron system; E3, cellular E3 ubiquitin ligase (CRBN in the case of dTAG-13). (B) Targeting strategy to achieve knock-in of FKBP-WRN at the endogenous human WRN locus. (C) Immunoblotting confirming successful knock-in of FKBP-WRN in human cancer cell lines treated or not with 0.5 μM dTAG-13 for 24 h. One of two independent experiments is shown. (D) Viability of FKBP-WRN expressing human cancer cell lines treated or not with 0.5 μM dTAG-13 for 6 days. Where indicated, shWRN expression was induced with 1 μg/mL doxycycline (DOX). Average (+/– S.D.) of three independent experiments is shown. (E) Immunoblotting comparing the kinetics of dTAG-induced FKBP-WRN degradation to that of shWRN-induced WRN depletion. One of two independent experiments is shown.

To validate the degron system, we administered a standard dose of 0.5 μM dTAG-13 to MSI-H cancer cells expressing FKBP-WRN, which engaged the cellular E3 ubiquitin ligase CRBN (Nabet *et al*., 2018) to induce WRN degradation (Fig 1C). Since our RKO and KM12 cells contained a pre-existing doxycycline-inducible shWRN (DOX-shWRN) cassette (van Wietmarschen *et al*., 2020), direct comparison could be readily made between dTAG– and shRNA-mediated WRN depletion. Importantly, we found that dTAG-13 treatment impaired the viability of FKBP-WRN expressing RKO and KM12 cells to a similar extent as DOX-shWRN, while their parental counterparts were eliminated by DOX-shWRN but not dTAG-13 (Fig 1D). Similarly, the MSI-H cell line HCT116, but not the MSS OVCAR8, were also highly sensitive to dTAG-13 treatment (Fig 1D), confirming the target specificity and MSI selectivity of our degron system. Detailed time course analysis in both MSI-H (RKO, KM12) and MSS (OVCAR8) cells revealed that approximately 90% of WRN protein was degraded by two hours, while depletion of WRN by DOX-shWRN was comparatively slower (Fig 1E). On the other hand, dTAG-13 did not affect the stability of unmodified WRN in parental lines (Fig 1C). Taken altogether, we concluded that our FKBP-WRN degron system can induce rapid and synchronous degradation of WRN protein, and therefore is a suitable tool to dissect how acute loss of WRN function is sensed and responded to by MSI-H cancer cells.

### Acute loss of WRN triggers immediate DNA damage in cycling cells

Previous studies using sh/siRNA and/or CRISPR have shown that DNA damage accumulates in MSI-H cancer cells several days after WRN ablation (van Wietmarschen *et al*., 2020) (Chan *et al*., 2019) (Lieb *et al*., 2019) (Kategaya *et al*., 2019). However, these studies lacked sufficient temporal resolution to answer when and how DNA damage was first produced following WRN depletion. Using our inducible degron system, we found that acute degradation of FKBP-WRN led to measurable increases in phosphorylated KAP1 (pKAP1), a marker of DNA double strand breaks, as soon as 2-4 h after the addition of dTAG-13 in RKO and KM12 cells (Fig 2A, B).

**Fig 2.**
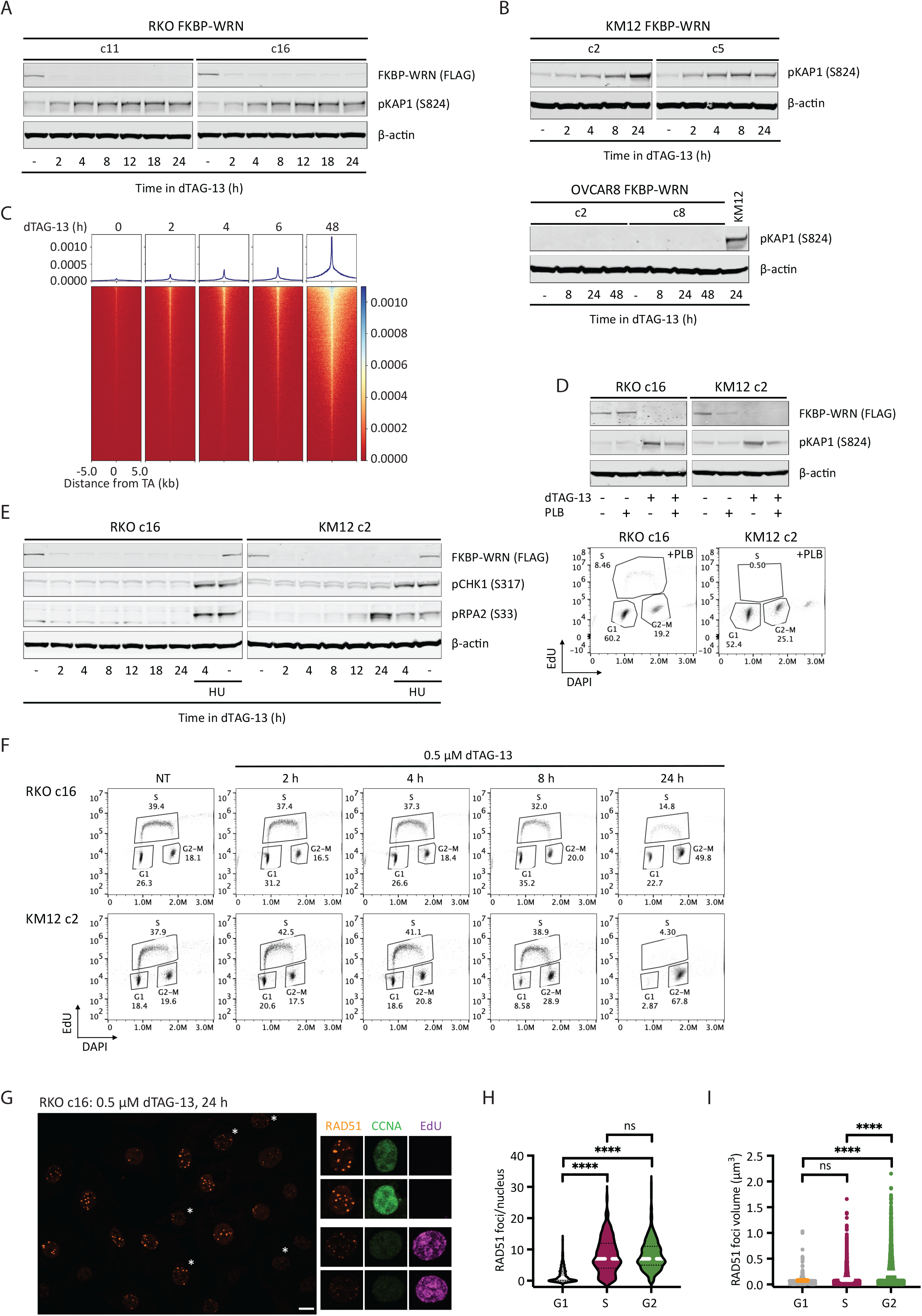
Acute degradation of WRN in MSI-H cancer cells causes replication-associated DNA damage without accompanying intra-S checkpoint activation. (A-B) Immunoblotting depicting the rapid induction of pKAP1, a marker of DNA double strand breaks, by 0.5 μM dTAG-13 in RKO (A) and KM12 (B) clones expressing FKBP-WRN. By contrast, dTAG-13 did not induce pKAP1 in OVCAR8 clones expressing FKBP-WRN even after extended treatment. One of two independent experiments is shown. (C) END-seq showing the rapid appearance of DNA breakage at unstable expanded TA repeats in KM12 cells treated with 0.5 μM dTAG-13. (D) Upper panel, immunoblotting showing that 0.5 μM dTAG-13 (24 h) induced significantly lower levels of pKAP1 in RKO and KM12 cells pre-treated for 24 h with 5 μM Palbociclib (PLB). Lower panel, successful inhibition of S-phase entry by PLB was confirmed by FACS–based cell cycle analyses. One of three independent experiments is shown. (E) Immunoblotting demonstrating the lack of CHK1 (S317) and RPA2 (S33) phosphorylation in dTAG-treated RKO FKBP-WRN cells. CHK1/RPA2 phosphorylation was also undetectable in KM12 FKBP-WRN cells except at 24 h after dTAG-13 treatment. Where indicated, cells were treated with 3 mM hydroxyurea (HU) for 2 h, either alone or in combination with 0.5 μM dTAG-13 (2 h dTAG-13 pre-treatment followed by 2 h HU+dTAG-13). Note that HU induced robust CHK1/RPA phosphorylation in both RKO and KM12 independently of WRN. One of two independent experiments is shown. (F) FACS-based cell cycle analyses showing that dTAG-treated RKO and KM12 cells failed to slow down or halt DNA synthesis during the first S-phase after acute WRN degradation. Note that WRN-degraded cells eventually completed S-phase and activated the G2 checkpoint. (G) Representative confocal images depicting RAD51 foci (maximum intensity projection) in RKO FKBP-WRN cells treated or not with 0.5 μM dTAG-13 for 24 h. Cyclin A (CCNA) and EdU positivity were used to distinguish cells in S-phase from those in G2. Note that RAD51 foci appear to be larger in G2 cells (EdU-negative, CCNA-high), as compared to S-phase cells (EdU-positive, CCNA-low/intermediate). (H) Quantification of RAD51 number per nucleus as a function of cell cycle position. (I) Volumetric analysis of RAD51 foci as a function of cell cycle position. One of two independent experiment is shown.

Consistent with this, breakage at sites of unstable TA repeats was also detected by ENDseq within hours of dTAG-13 treatment in KM12 (Fig 2C). Thus, MSI-H cells appeared to incur DNA damage almost immediately after FKBP-WRN has been degraded. By contrast, MSS OVCAR8 cells did not show induction of pKAP1 in response to WRN degradation, even days after dTAG-13 treatment (Fig 2B). Next, we examined whether DNA damage induction upon acute WRN loss was dependent on cell cycle position. Notably, while dTAG-induced WRN degradation itself was cell cycle-independent, blocking S phase entry with palbociclib (PLB) significantly reduced pKAP1 induction in RKO and KM12 cells as compared to their asynchronous counterparts (Fig 2D). These observations suggested that DNA damage caused by loss of WRN depends on active proliferation.

### WRN deficient in MSI-H cells mount a non-canonical DDR

Eukaryotic cells typically respond to DNA damage by arresting cell cycle progression through the prompt activation of checkpoints, orchestrated by ATM-CHK2 and ATR-CHK1 (Waterman Haber & Smolka, 2020). In WRN-degraded MSI cancer cells, we found that the rapid elevation of pKAP1 was closely followed by the induction of CHK2 phosphorylation (Appendix Fig S2A). Therefore, the presence of DNA DSBs was clearly sensed by ATM. However, WRN degradation triggered little to no CHK1 phosphorylation, although the ATR-CHK1 pathway was intact in these cells and could be activated by hydroxyurea (Fig 2E and Appendix Fig S2B).

In the absence of CHK1-enforced intra-S checkpoint, DNA replication proceeded unperturbed in WRN-degraded RKO and KM12 cells for at least 8 h (Fig 2F). By 24 h, we detected a large accumulation of cells with 4N DNA content, indicating that most cells have completed S-phase and entered G2, where CHK2 activity had presumably slowed down their progression towards mitosis. Interestingly, these G2 cells contained numerous prominent RAD51 foci that appeared qualitatively larger and brighter than RAD51 foci typically found in S-phase cells (Fig 2G, H, Appendix Fig S2C). Detailed volumetric analyses subsequently confirmed that RAD51 foci present in G2 were indeed larger than those found in S-phase (Fig 2I). By contrast, ionizing irradiation induced RAD51 foci were of similar sizes irrespective of cell cycle position in S or G2 (Appendix Fig S2C). Although the exact significance of this difference remains unclear, we speculate that the lack of robust checkpoint early on may limit the repair of replication-associated DNA damage in WRN deficient MSI-H cancer cells, and as such continuously concentrate repair factors at DNA damage sites.

### WRN-degraded MSI-H cancer cells propagate and amplify DNA damage across multiple mitoses

To gain additional insight into how MSI-H cancer cells respond to WRN ablation, we constructed an RKO FKBP-WRN cell line suitable for live cell imaging (Fig 3A, B). This system combines three separate reporters, enabling us to monitor chromatin (H2B-mTurquoise), cell cycle (PIP-NLS-mVenus) and DNA damage (53BP1-mCherry) simultaneously and in real-time. We first confirmed that acute degradation of WRN indeed induced DNA damage, marked by 53BP1, very rapidly (Fig 3C, D) and preferentially in S-G2 phases of the first cell cycle (Fig 3D, E). The behavior of these 53BP1 foci was quite heterogeneous, with some being highly dynamic while others were relatively persistent. In accordance with previous studies (Giunta Belotserkovskaya & Jackson, 2010; Nelson Buhmann & von Zglinicki, 2009), we observed that all 53BP1 foci dissolved in early mitosis. Nevertheless, we found that more than half of WRN-degraded cells harbored unrepaired DNA breaks (chromatid– and/or chromosome-type) in the first metaphase, and 10% additionally displayed complex rearrangements that were indicative of erroneous repair (Fig 3F). Thus, cellular DNA repair machineries were not able to completely heal all WRN-associated lesions prior to cell division. Consistent with this idea, most WRN-degraded cells appeared to regain 53BP1 foci in the following G1 and the median 53BP1 foci per nucleus rose steadily over time (Fig 3C).

**Fig 3.**
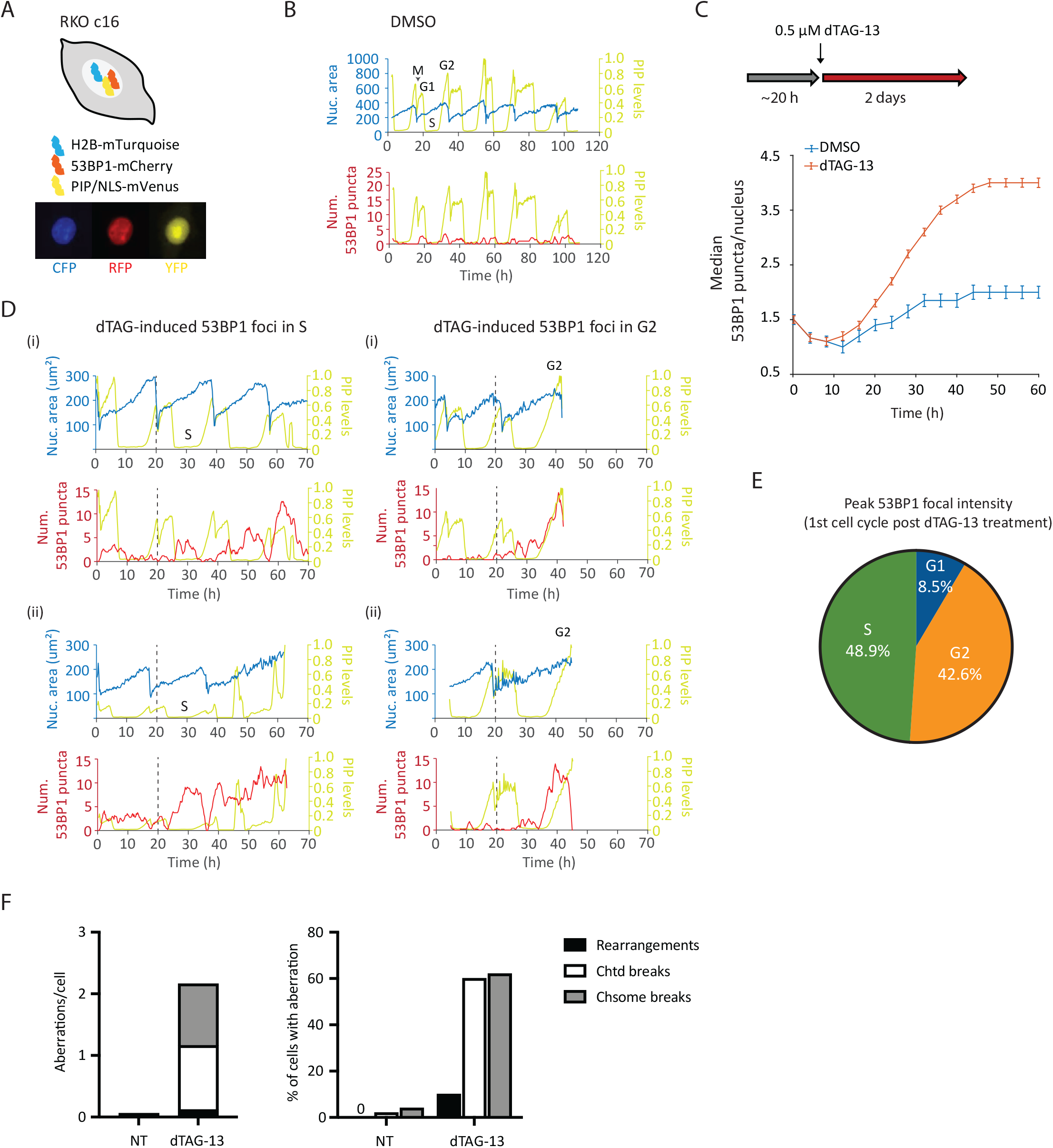
Induction and repair of DNA damage in WRN-degraded MSI-H cancer cells monitored by time-lapse microscopy. (A) Schematics of the RKO triple reporter line used for live cell imaging. An example of a vehicle-treated cells is shown below in the CFP, RFP and YFP channels. (B) Automated cell tracking was performed as described in the Methods section. Example traces of a vehicle-treated reporter cell over time; different phases of a cell cycle are indicated. The birth of a cell at mitosis (M) coincides with a reduction in nuclear area (as defined by H2B-mTurquoise) as well as PIP sensor intensity. PIP intensity increases in G1 but is then abruptly lost as a cell enters S-phase. Finally, PIP expression returns in G2. 53BP1 foci formation is also shown for this vehicle-treated cell. (C) Formation of 53BP1 puncta as a function of time. Cells were first imaged for approximately 20 h without drug to establish baseline 53BP1 levels. Subsequent addition of dTAG-13 resulted in a rapid induction of 53BP1 puncta over the baseline, which steadily increased over time. One of two independent experiments is shown. (D) Example traces of four dTAG-treated reporter cells. The two cells on the left panels showed induction of 53BP1 puncta during the first S-phase after WRN degradation. The two cells on the right panels had primarily induced 53BP1 in the first G2 after WRN degradation. Note however that these two cells also displayed a smaller induction of 53BP1 in the preceding S-phase. (E) Quantification of 53BP1 puncta intensity during the first cell cycle following acute WRN degradation revealed that peak 53BP1 induction occur predominantly in S and G2. One of two independent experiments is shown. (F) Analyses of metaphase spreads showed that a majority of dTAG-treated RKO cells failed to completely repair DNA breaks incurred in S/G2 and carried residual damages into the first mitosis. One of two independent experiments is shown.

To examine how transmission of unresolved DNA damage may impact future cell division activities, we manually traced the lineage of a subset of RKO reporter cells over a span of six days. Overall, we found that most of vehicle-treated RKO cells (n=25) divided at least 6 times (after which accurately tracking became difficult due to cell movements and crowding), while WRN-degraded RKO cells (n=50) divided on average 2-3 times (range 0-6) (Fig 4A).

**Fig 4.**
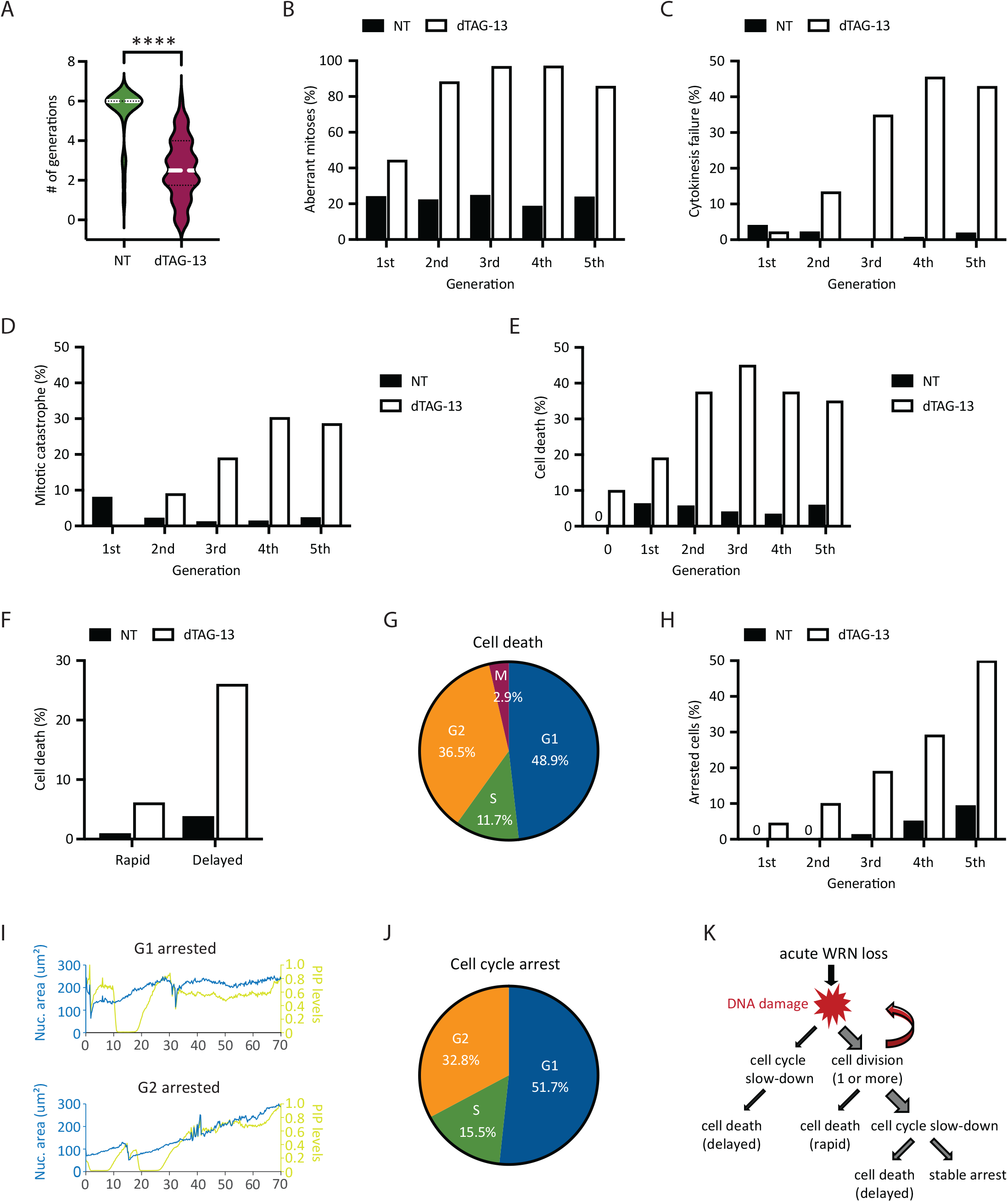
Heterogeneity in the fate of WRN-degraded MSI-H cancer cells. RKO reporter cells were treated or not with 0.5 μM dTAG-13 and imaged by time-lapse microscopy over a span of 6 days. Manual tracking was performed on 25 vehicle-treated and 50 dTAG-treated cells. (A) Cell division activities. Most of vehicle-treated cells divided at least 6 times, producing 840 tracked progeny. By contrast, dTAG-treated cells divided less frequently (average 2-3 times, range 0-6) and produced less progeny (377 tracked). (B) Mitotic abnormalities as a function of cell division. A mitosis was scored as abnormal if it exhibited one or more of the following features: bridging, micronuclei formation, cytokinesis failure and/or mitotic catastrophe. (C) Frequency of cytokinesis failure as a function of cell division. (D) Frequency of mitotic catastrophe as a function of cell division. Mitotic catastrophe was defined as any mitosis that resulted in complete failure to divide (failed mitosis), multipolar chromosome segregation or rapid induction of mitotic cell death. (E) Frequency of cell death as a function of cell division. Note that 10% of dTAG-treated cells died without undergoing a single mitosis (denoted as generation 0). (F) Cell death as a function of time. A death event is defined as rapid if it occurred <10 h after the latest mitosis from which the cell was born. Otherwise, it is defined as delayed. (G) Cell death as a function of cell cycle position. (H) Frequency of cell cycle arrest as a function of cell division. (I) Example traces of two dTAG-treated cells showing arrest in G1 (upper panel) or G2 (lower panel). (J) Cell cycle arrest as a function of cell cycle position. One of two independent experiments is shown. (K) Diagram depicting the various observed fates of RKO cells following acute WRN degradation.

Approximately 45% of first mitoses in dTAG-treated cells exhibited discernible abnormalities (Fig 4B), such as chromosomal bridges and micronucleation. Subsequent mitoses in their progeny were even more error-prone, causing severe defects such as cytokinesis failures and mitotic catastrophe, in addition to bridges and micronuclei (Fig 4B-D). By contrast, the frequencies of abnormal mitoses among vehicle-treated cells were significantly lower and stayed constant with each generation (Fig 4B-D). Altogether, these observations support a model wherein WRN deficient MSI-H cells propagate and amplify DNA damage in a trans cell cycle manner due to a combination of inadequate repair and accrual of new lesions. As such, the massive genome fragility we and others have previously detected in MSI-H cancer cells after several days of prolonged WRN depletion via RNA interference likely reflects the culmination of DNA damage over multiple cell cycles rather than a single catastrophic event.

### WRN-degraded MSI-H cancer cells exhibit marked heterogeneity in their fate

Previously, it was reported that MSI-H cancer cells in which WRN was depleted by si/shRNA ultimately died in a manner consistent with induction of apoptosis (Chan *et al*., 2019; Kategaya *et al*., 2019). To validate and extend these findings, we conducted detailed manual cell fate mapping using our live imaging reporter system. We observed that 10% of dTAG-treated RKO cells died without ever dividing, whereas none of the vehicle-treated cells did so (Fig 4E). Of those dTAG-treated cells that divided at least once, we carefully documented the fate of 377 progeny, of which 35% died within six days. By contrast, 4.9% of their vehicle-treated counterparts (840 tracked progeny) died during the same period. In concordance with the trans cell cycle propagation and amplification of DNA damage, we found that death following acute WRN loss tended to occur after at least two rounds of cell division, while the rate of death among vehicle-treated cells remained constant over time (Fig 4E). Interestingly, the time elapsed between the birth of a given cell at mitosis and its subsequent demise was typically quite protracted (>10 h up to days) (Fig 4F). During this time, the doomed cell might linger in the ensuing G1, attempt to replicate its DNA, or even enter G2 after completion of S-phase (Fig 4G). However, a minority of cells died rapidly (<10 h) after birth, either while still in M or shortly thereafter in G1 (Fig 4F). Most of the observed cell death events exhibited chromatin condensation, consistent with the induction of apoptosis. However, we noted that in many cases, the nucleus also seemed to “explode”, while in other instances the nucleus remained relatively intact (Appendix Movie). Overall, these data suggested that WRN-deficiency triggers cell death with significant temporal and morphological heterogeneity in MSI-H cancer cells, perhaps involving mechanisms in addition to classical apoptosis.

Of the cells that were still alive by the end of the experiment, we found that a significant proportion (15-20% of all tracked progeny) appeared to have entered a state of stable arrest. As with cell death, the likelihood of cell cycle arrest also increased with each division (Fig 4H). Similarly, CFSE dye dilution assay showed a progressive decline in cell division activities for RKO, KM12 and HCT116 cells following acute WRN degradation (Appendix Fig S3A), mirroring a steady rise in the levels of phosphorylated CHK2 over time (Appendix Fig S3B, C). We found that CHK1 activation remained undetectable in RKO FKBP-WRN clones even at late time points (Appendix Fig S3B), and accordingly, careful manual tracking of WRN-degraded RKO reporter cells showed that they arrested predominantly in G1 or G2, with only a minor fraction locked in S-phase (Fig 4I, J). Interestingly, WRN-degraded KM12 and HCT116 cells were able to activate CHK1 at late time points in addition to CHK2, which may explain why dTAG-13 treatment caused a stronger proliferative impairment in these two cell lines compared to RKO (Appendix Fig S3A). Altogether, these data highlight the heterogeneous fate that acute WRN loss imparts on MSI-H cancer cells (Fig 4K).

### MSI-H cancer cells do not readily adapt to WRN deficiency

Given that WRN addiction is a striking and potentially actionable vulnerability in human MSI-H cancers, a key question that has not yet been addressed is whether WRN hypersensitivity can circumvented by secondary inactivation of another cellular process. To test this possibility, we conducted a knockout CRISPR screen in a clone of Cas9-expressing RKO cells by transducing them with a genome wide lentiviral library (Brunello) of single-guide RNAs (sgRNAs). The resultant pools of edited cells were either treated with vehicle or exposed to a near-lethal dose of dTAG-13 over 6 days (Fig 5A). We used the model-based analysis of genome-wide CRISPR-Cas9 knockout (MAGeCK) algorithm to calculate gene enrichment or depletion (Li *et al*, 2014). Importantly, we identified CRBN, the ubiquitin E3 ligase engaged by dTAG-13, as the top hit whose deletion confers resistance to dTAG-13 (Fig 5B), thereby confirming the validity of our approach.

**Fig 5.**
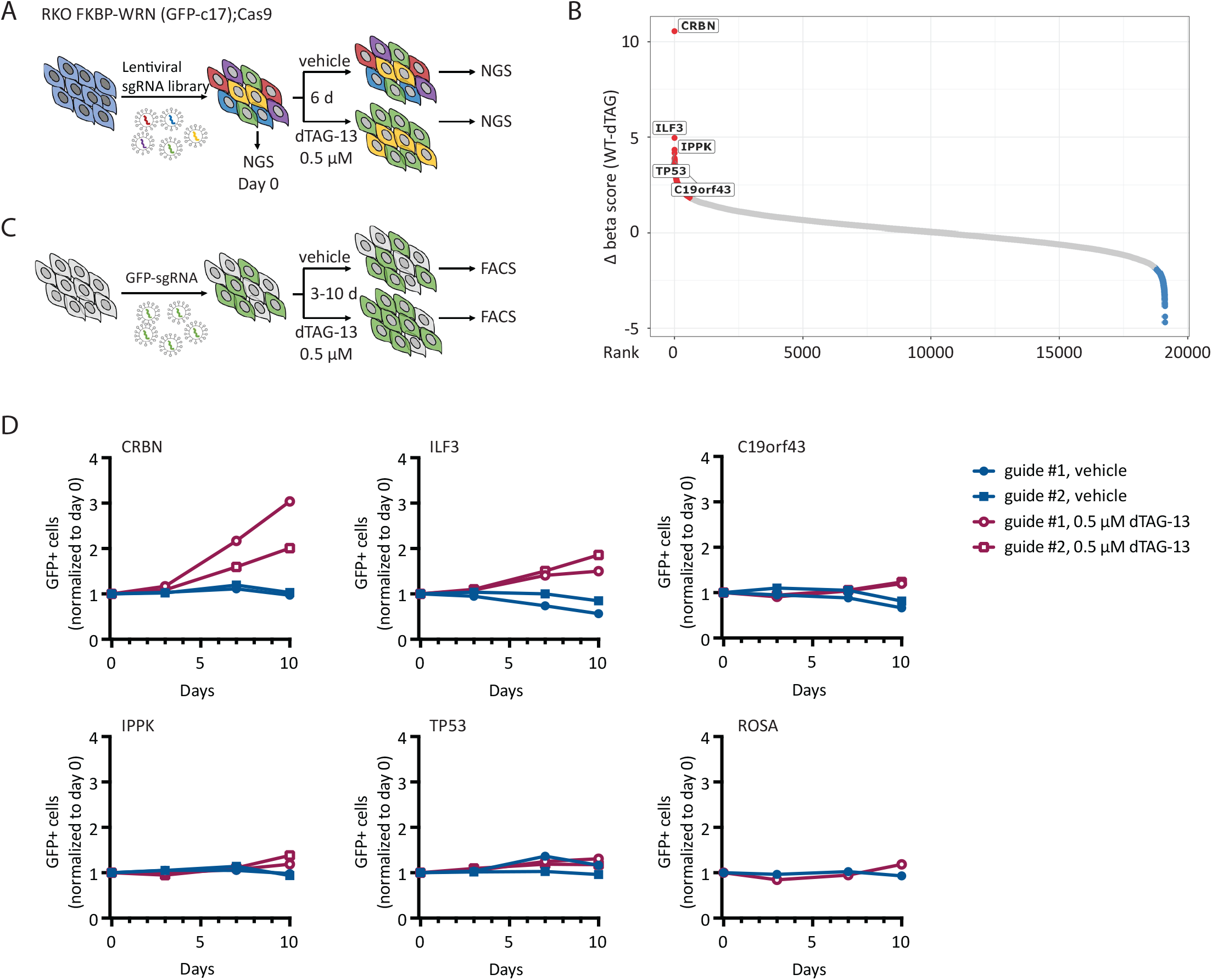
MSI-H cancer cells do not readily adapt to WRN deficiency. (A) Schematics of the CRISPR screening strategy used in RKO c17/Cas9 cells to identify genetic hits that could potentially confer resistance to WRN loss. (B) Candidate resistance genes identified in the screen. (C) Schematics of growth competition assays used to validate candidate resistance genes. (D) Growth competition assays confirmed that deletion of CRBN, and to a lesser degree ILF3, rescued cytotoxicity caused by acute loss of WRN in RKO c17/Cas9 cells. Note that CRBN encodes the cellular E3 ubiquitin ligase that is engaged by dTAG-13 to promote FKBP-WRN degradation. One of two independent experiments is shown.

Apart from CRBN, we identified only a few additional candidate resistance genes, including ILF3, IPPK, TP53 and c19orf43 (Fig 5B). Nevertheless, the sgRNAs targeting these genes showed much lower levels of enrichment than guides targeting CRBN. To validate potential hits, we transduced the RKO clone used in the original CRISPR screen with two individual guides targeting these candidate genes and performed growth competition assays in the presence or absence of dTAG-13 (Fig 5C). We confirmed that depletion of CRBN completely abrogated dTAG-induced FKBP-WRN degradation and as expected, CRBN-deficient RKO cells were significantly enriched after dTAG-13 treatment (Fig 5D, Appendix Fig S4A, B). Depletion of ILF3 also conferred some level of resistance to dTAG-13 (Fig 5D); however, cells deficient in ILF3 no longer appeared to degrade FKBP-WRN efficiently in response to dTAG-13 treatment (Appendix Fig S4B), thus ruling it out as a *bona fide* WRN resistance gene. Finally, the loss of IPPK, c19orf43 or TP53 had modest to negligible impact on cell viability under conditions of WRN deficiency (Fig 5D). These findings were recapitulated in a different clone of RKO and in KM12 (Appendix Fig S4C, D). Overall, we conclude that there is currently no evidence of any generalized mechanism of WRN resistance in human MSI-H cancer cells.

### ATR inhibition sensitizes MSI-H cancer cells to partial loss of WRN

The apparent inability of human MSI-H cancer cells to adapt to WRN deficiency is highly encouraging and supports the therapeutic utility of WRN inhibitors. Clinically, it is often impossible to completely inactivate a target protein *in vivo* because of excessive toxicity and/or limited bioavailability. When we lowered the dose of dTAG-13 from 0.5 μM to the 2.5-10 nM range, we observed both higher levels of residual WRN protein and increased cell survival (Appendix Fig S5A-C), suggesting that low doses of dTAG-13 treatment can be used to simulate partial inhibition of WRN.

Next, we explored ways to augment the efficacy of low dose dTAG-13 treatment. Previous studies from our lab have shown that WRN phosphorylation by ATR is required for its genome protective functions in MSI-H cancer cells (van Wietmarschen *et al*., 2020). We therefore tested whether ATR inhibition could complement low dose dTAG-13 treatment, potentially by impairing the function of residual active WRN. However, we found that MSI-H cancer cells were highly sensitive to the ATR inhibitor AZ-20 even under WRN-proficient conditions, especially at doses of 0.5 μM and above (Appendix Fig S5D, E and data not shown). This notion was further supported by data extracted from several genome-wide screens and public drug sensitivity datasets (Appendix Fig S5F, G). While these results support the potential use of ATR inhibitor monotherapy in MSI-H human cancers, the inherent hypersensitivity of MSI-H cells to ATR inhibitors made it difficult to interpret combinatorial effect of dual WRN/ATR inhibition. To circumvent this issue, we decided to combine low doses of dTAG-13 with concentrations of AZ-20 that were lower than those used in our single agent studies. Notably, we found that the combination regimen imparted significantly greater losses in the viability of RKO, KM12 and HCT116 cells than what was achievable with either agent alone (Fig 6A-C and Appendix Fig S6A, B). On the contrary, the combination regimen yielded comparable effect as high dose ATR inhibition alone in MSS OVCAR8 cells (Fig 6D). Mechanistically, the combination regimen led to increased DNA damage load, as shown by elevated KAP1 phosphorylation (Fig 6E) and breakage at TA repeat sites (Appendix Fig S6C). Corroborating these findings, live cell imaging showed that 53BP1 foci formation was significantly enhanced in RKO reporter cells treated with the combination regimen as compared to their single agent treated counterparts (Fig 6F), which likely contributed to decreased cell proliferation, augmented cell cycle arrest and more rapid induction of cell death (Fig 6G-I). Taken together, these data provide a compelling rationale for the further development and testing of combination regimens of ATR and WRN inhibitors to treat human MSI-H cancers.

**Fig 6.**
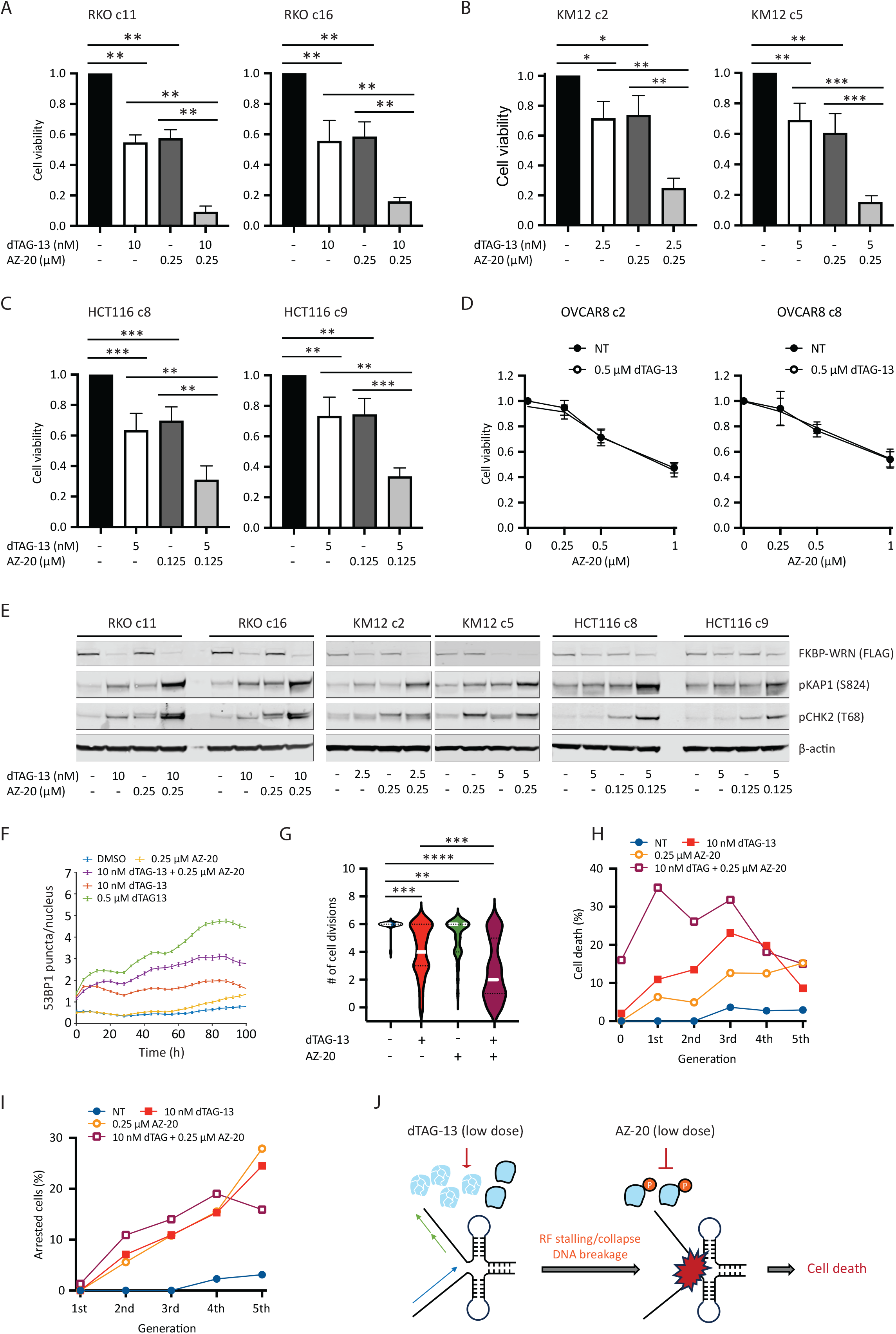
ATR inhibition can compensate for incomplete WRN degradation. (A-C) Human cancer cells expressing FKBP-WRN were treated as indicated for 6 days. When combined, the ATR inhibitor AZ-20 was always added two hours after the start of dTAG-13 treatment. Viability assays showed that the combination regimen produced significantly superior cytotoxicity as compared to single-agent treatments in MSI-H RKO (A), KM12 (B) and HCT116 (C) clones, but not in MSS OVCAR8 (D). Average (+/– S.D.) of three (OVCAR8, KM12 c2) or four (RKO, HCT116, KM12 c5) independent experiments is shown. (E) MSI-H cancer cells expressing FKBP-WRN were treated as indicated for 24 hours. Immunoblotting showing that the combination regimen resulted in markedly elevated levels of pKAP1 as compared to either single agent alone. One of two independent experiments is shown. (F) RKO reporter cells were treated as indicated and subjected to live cell imaging. Automated tracking of 53BP1 puncta showed that the combination regimen induced more 53BP1 than either agent alone. (G-I) Manual fate mapping of RKO reporter cells: vehicle (n=25, 1094 tracked progeny), 10 nM dTAG-13 (n=50, 992 tracked progeny), 0.25 μM AZ-20 (n=40, 1074 tracked progeny), dTAG-13/AZ-20 (n=50, 497 tracked progeny). Cells treated with the combination regimen exhibited reduced cell division activities (G) accompanied by increases in both cell death (H) and cell cycle arrest (I). For (F-I), one of two independent experiments is shown. (J) Proposed model of how ATR inhibition compensates for incomplete WRN degradation.

## Discussion

Successful targeting of tumor-specific vulnerabilities is the ultimate goal of modern precision anti-cancer therapy. Recently, the RECQ helicase WRN has emerged as a crucial survival factor in human MMR-deficient MSI-H cancers (Morales-Juarez & Jackson, 2022) (Picco *et al*., 2021) (van Wietmarschen *et al*., 2020) (Chan *et al*., 2019) (Behan *et al*., 2019) (Lieb *et al*., 2019) (Kategaya *et al*., 2019). The unique specificity of WRN addiction in MSI-H cancer cells and the extreme toxicity unleashed by WRN loss puts it on par with PARP inhibition in BRCA1/2-deficient tumors (Morales-Juarez & Jackson, 2022) (van Wietmarschen Nathan & Nussenzweig, 2021). Despite holding considerable promise, however, clinical grade WRN inhibitors are not currently available. To get a sense of how MSI-H cancer cells might respond to pharmacological WRN inactivation in both the short and long term, we engineered a rapidly inducible degron system and probed it with state-of-the art reporter-based live cell imaging. We discovered that accrual of replication-associated DNA double strand breakage is a conserved immediately early consequence of acute WRN loss, while cell death and cell cycle arrest occur relatively late in a more heterogeneous manner.

The lesions that accumulate in the genomes of WRN-degraded MSI-H cancer cells are quickly sensed by ATM, leading to the mobilization of many canonical signaling and repair factors including pKAP1, pCHK2, 53BP1, RPA and RAD51. Surprisingly, however, MSI-H cancer cells fail to robustly activate CHK1 during the first S-phase after acute WRN loss, thus allowing replication to continue unabated and eventually be completed in the presence of DNA damage. Through real-time monitoring of 53BP1 foci, we also found that a substantial fraction of WRN-degraded RKO cells appears to incur additional DNA damage in G2 (Fig 3D), possibly the result of nucleolytic processing of unresolved aberrant replication intermediates. In this manner, the lack of a viable intra-S checkpoint early on could trigger a cascade of damage accumulation as cells progress through S/G2, which ultimately overwhelms the cellular DNA repair machineries. Moreover, without activated CHK1 to support pCHK2, the G2 checkpoint is likely to become more vulnerable to slippage. Indeed, examination of metaphase chromosomes of RKO cells entering the first mitosis after WRN degradation revealed both unresolved DNA breaks and genomic rearrangements associated with mis-repair. Thus, the inability of MSI-H cancer cells to halt DNA synthesis immediately upon WRN loss leads to transmission of DNA damage to the next cell cycle, which could be a major cause of subsequent cytotoxicity. We further suggest that these effects should be recapitulated by acute pharmacological inhibition of WRN.

By carefully tracking the fate of WRN-degraded RKO reporter cells, we found that they typically progress through two or more cell cycles before succumbing. In addition, some cells apparently entered a particular cell cycle phase from which they did not emerge, although it is not yet clear whether this leads to a permanent exit from the cell cycle (i.e., senescence), or if it’s simply a prolonged transitional state after which cells either die or recover. Previously, it was reported that WRN-deficient MSI-H cancer cells undergo apoptosis, a form of cell death that is non-immunogenic. However, based on morphological characteristics and a relative lack of PARP1 cleavage (not shown), our data raises the possibility that cell death triggered by WRN deficiency in MSI-H cancer cells may be biochemically distinct from classical apoptosis. If WRN inhibition indeed induces potentially immunogenic forms of cell death as well as senescence in MSI-H tumors, combining it with immunotherapy may prove to be even more beneficial.

As with any targeted therapy, the clinical benefit of prospective WRN inhibitors is potentially limited by the emergence of resistance mechanisms. In this regard, a recent paper reported that inactivation of TP53 completely abrogated apoptosis induced by WRN depletion in MSI-H colorectal cancer cells (Hao *et al*, 2022). However, TP53 did not behave as a *bona fide* WRN resistance gene in our system. In addition, KM12 cells are known to be p53-mutated yet responded strongly to WRN depletion by either dTAG-13 or shRNA. Moreover, both we and others (Lieb *et al*., 2019) have observed that p53-deficient HCT116 cells were just as (hyper)sensitive as their WT counterparts to si/shWRN (not shown). Overall, aside from a few potential mild and clone-specific modifiers, we found no evidence of any generalized mechanisms of that could confer resistance to WRN deficiency in MSI-H cancer cells. Nevertheless, we cannot rule out the possibility that such resistance might be obtained through overexpression of certain cellular factors and/or gain-of-function mutations. In addition, we anticipate that mutations altering the binding surfaces of prospective WRN inhibitors could potentially drive clinical resistance in a manner described previously for PARP inhibitors (Pettitt *et al*, 2018).

Finally, just how much residual WRN is sufficient to promote survival of MSI-H cancers is a key question that should be considered when developing future WRN inhibitors. Using our FKBP-WRN degron system, we observed the most efficient cell killing at doses of dTAG-13 (0.04-0.5 μM) that degraded approximately 90% of WRN protein, while dTAG-13 doses below this range resulted in significantly less cytotoxicity. Because this applied to both homozygous and hemizygous clones, we estimated that about 95% of functional WRN protein would need to be inhibited for optimal outcome. In other words, very little WRN may be able to sustain the viability of MSI-H cancer cells. Interestingly, previous findings have shown WRN to be functionally regulated by ATR-mediated phosphorylation (van Wietmarschen *et al*., 2020) (Ammazzalorso *et al*, 2010) (Pichierri Rosselli & Franchitto, 2003), suggesting a combination approach might promote stronger suppression of WRN activity. Indeed, we were able to significantly enhance cytotoxicity in MSI-H cancer cells by combining low doses of dTAG-13 with relatively low concentrations of the ATR inhibitor AZ-20, with the efficacy of the combination regimen approaching that of high dose dTAG-13. We hypothesize that the combination works by augmenting partial degradation of WRN with functional inactivation of the remaining proteins, effectively achieving near complete inhibition of WRN (Fig 6J). The inhibition of ATR may additionally subvert a critical genome surveillance mechanism that normally promotes RF stability and repair, compounding the damage leading to replication catastrophe (Fig 6J). Whatever the precise mechanism, our data provided a compelling rationale to further explore productive combination regimens of ATR and WRN inhibitors for the treatment of human MSI cancers.

## Materials and Methods

### Cell lines and culture conditions

RKO, KM12, SW48, NCIH508, SW620 and SW837 cells containing doxycycline-inducible shWRN cassette were generated as previously described (Chan *et al*., 2019; van Wietmarschen *et al*., 2020). HCT116, LS180 and OVCAR8 cells were obtained from commercial sources. RKO and LS180 were grown under standard conditions in EMEM (ATCC). KM12, SW48, NCIH508, SW837 and OVCAR8 were cultured in RPMI-1640, while HCT116 and SW620 were maintained in McCoy 5A and Leibovitz L-15 media, respectively. RPMI-1640, McCoy 5A and L-15 were from Thermo Fisher Scientific. All media were supplemented with 10% heat-inactivated fetal bovine serum (FBS, Gemini Bio-Products) and 1% antibiotic (penicillin and streptomycin, GIBCO). All cell lines were confirmed free of mycoplasma.

### Construction of FKBP-WRN donor and WRN targeting vector

DNA encoding the 5’– and 3’-homology arms of WRN was obtained from Twist Biosciences and subsequently cloned into pGMC00018, along with the PCR product encoding Puromycin^R^-2A-3X-FLAG-FKBP12 or eGFP-2A-3X-FLAG-FKBP12, using isothermal assembly. Guide RNAs targeting the N-terminus of WRN (ENST00000298139.6) were designed using sgRNA Scorer 2.0 (Chari *et al*, 2017). Candidates 3537 (TGACACCTAGGTCCAAGCAT**AGG,** PAM in bold) and 3539 (TGGAAACAACTGCACAGCAG**CGG**) were chosen based on superior cutting efficiency in initial test experiments. Oligonucleotides corresponding to these two guide sequences were subsequently annealed and ligated into the Cas9-containing pDG458 vector (Adikusuma Pfitzner & Thomas, 2017) using golden gate assembly; pDG458 was a gift from Dr. Paul Thomas (Addgene plasmid #100900, RRID:Addgene_100900). The FKBP-WRN donor plasmid construct and WRN targeting vector were verified using Sanger sequencing (ACGT DNA Sequencing Services).

### Generation of FKBP-WRN cell lines

Recipient cells were transiently transfected with both the FKBP-WRN donor and the WRN targeting vector using X-tremeGENE 9 (Sigma), as per manufacturers’ instructions. After 48 h, cells were either subjected to puromycin selection (if Puromycin^R^-2A-3X-FLAG-FKBP12 was used as the donor, as in RKO and KM12) or FACS sorted (if eGFP-2A-3X-FLAG-FKBP12 was used as the donor, as in HCT116 and RKO c17). Single cell derived candidate clones were harvested for DNA and protein. The targeted region was amplified by PCR (MyTaq Red mix, Bioline) from 100 ng of genomic DNA. Successful knock-in of FKBP-WRN was confirmed by Sanger sequencing and by immunoblotting (for both WRN and FLAG).

To generate the FKBP-WRN triple reporter cell line, RKO clone #16 was stably transduced with lentiviral vectors encoding H2B-mTurquoise, PIP/NLS-mVenus and 53BP1-mCherry. H2B-mTurquoise (Spencer *et al*, 2013) and PIP/NLS-mVenus (Grant *et al*, 2018) have been described previously, cell cycle (PIP-NLS-mVenus). 53BP1-mCherry was excised mCherry-BP1-2 pLPC-Puro (Addgene plasmid #19835) (Dimitrova *et al*, 2008) and inserted into H53-LV (gift of Dr. Michael Ward) using isothermal assembly. Triple-positive cells were FACS-sorted (Sony MA900) and expanded for future use.

### Chemicals

dTAG-13 was purchased from Tocris Bioscience. Doxycycline, hydroxyurea and CFSE (carboxyfluorescein diacetate N-succinimidyl ester) were from Sigma. Palbociclib (PLB) and AZ-20 were from Selleck Chemicals. EdU (5-ethynyl-2’-deoxyuridine) was from Invitrogen. Formaldehyde and paraformaldehyde were obtained from Sigma and Electron Microscopy Sciences, respectively. DAPI (4′,6-diamidino-2-phenylindole) was from Thermo Fisher Scientific. All other chemical reagents were from Sigma.

### Cell viability assays

Cells were seeded in 6-well plates (10,000-20,000 per well) overnight and treated with drugs the following day. DOX-shWRN expression was induced with 1 μg/mL doxycycline. The standard dosage of both dTAG-13 and AZ-20 was 0.5 μM, with DMSO serving as the vehicle. Where indicated, lower doses of either or both were also used. When used in combination, AZ-20 was always added 2 h after dTAG-13. Cells were treated continuously for 6 days. Viability was determined using the CellTiter-Glo Luminescent Cell Viability Assay (Promega) as per manufacturer’s instructions.

### Live cell imaging

RKO FKBP-WRN triple reporter cells were seeded in a 24-well plate (Ibidi, #89626) approximately 18 hours prior to imaging. Cells were plated such that cell density remained sub-confluent until the end of the imaging period (3-6 days). Depending on the goal of the experiment, different drug treatment and imaging schedules were used. To examine 53BP1 induction, cells were first imaged for approximately 20 h in drug-free media to establish baseline 53BP1 level and behavior, after which 0.5 μM dTAG-13 or vehicle (DMSO) was quickly added and imaging resumed for an additional three days. To conduct long-term cell fate mapping, cells were treated with DMSO or 0.5 μM dTAG-13 and imaged over a span of six days. To explore the effect of WRN/ATR single or co-inhibition, cells were first treated with vehicle or 10 nM dTAG-13, followed by the addition of vehicle or 0.25 μM AZ-20 two hours later before commencing image acquisition (6 days). In all cases, time-lapse images were taken in CFP, YFP, and RFP channels every 12 min on a Nikon Ti2-E inverted microscope (Nikon) with a 20X 0.45NA objective. NIS Elements (Nikon, v5.11.00) software was used for controlling image acquisition. Total light exposure time was kept under 600 msec for each time point. The microscope was housed within an environmental chamber to maintain cells at 37 °C with 5% humidified CO_2_.

### Image processing and single cell tracking

All image analyses were performed with custom MATLAB scripts as previously described (Cappell *et al*, 2016). Cells were segmented for their nuclei based on H2B-mTurquoise. Nuclear PIP sensor activity was calculated as median nuclear intensity and then normalized to obtain a range of values between 0 and 1. 53BP1-mCherry puncta corresponding to pixel locations within the nuclear mask were quantified by applying a Top-hat filter followed by segmentation. The number of unique segmented objects corresponding to the mCherry channel were counted to obtain the number of 53BP1 puncta, and the pixel intensity of each spot was quantified to obtain 53BP1 puncta intensity.

Segmentation and automated tracking of cells from time-lapse microscopy films was performed using a previously published MATLAB pipeline (Cappell *et al*., 2016); code available at https://github.com/scappell/Cell_tracking. Briefly, cell tracks were linked by screening the nearest future neighbor using segmented cell nuclei in adjacent frames. When the imaging plate was removed from and put back on the microscope stage (for addition of drugs), the plate jitter was calculated by registering images of the nucleus-stained channel and corrected prior to tracking.

Fate mapping of a subset of cells was performed by manual analyses of time-lapse microscopy films using Fiji (Schindelin *et al*, 2012). Only cells that were present in the first time-lapse frame and not already showing overt signs of abnormalities were included in the analyses. Care was also taken to exclude cells that rapidly moved out of frame. Cell cycle positions were assigned based on manual examination of PIP sensor intensity.

### Immunoblotting

Western blotting was performed as previously described (Zong *et al*, 2015). Briefly, cells were washed in PBS and lysed in an extraction buffer containing 50 mM Tris-HCl (pH 7.5), 200 mM NaCl, 5% Tween-20, 2% Igepal, 2 mM PMSF, 2.5 mM b-glycerophosphate (all from Sigma) and supplemented with protease and phosphatase inhibitor tablets (both Roche Diagnostics). Equal amounts of protein were loaded into precast 4-12% Bis-Tris mini-gels (Invitrogen) and resolved by SDS-PAGE. Thereafter, proteins were transferred onto nitrocellulose membranes, blocked in a TBS-based blocking agent (LiCor Biosciences) and incubated with the corresponding primary antibodies recognizing the following proteins: WRN (1:1000, Novus), FLAG (1:1000, Sigma), pKAP1 (S824, 1:1000, Bethyl), pATR (T1989, 1:500, Cell Signaling), pCHK1 (S345, 1:500, Cell Signaling), pCHK1 (S317, 1:500, Novus), pCHK1 (S296, 1:1000, gift of Dr. Lee Jung-Min), CHK1 (1:1000, Cell Signaling), pRPA2 (S33, 1:1000, Bethyl), RPA2 (1:2000, Cell Signaling), pCHK2 (T68, 1:1000, Cell Signaling), CHK2 (1:1000, Cell Signaling), β-actin (1:5000, Sigma) and Tubulin (1:5000, Sigma). Fluorescently labeled secondary antibodies were used at a dilution of 1:10,000 (Li-Cor Biosciences). Visualization and quantification of protein bands was achieved by fluorescence imaging using a Li-Cor Odyssey CLx imaging system (Li-Cor Biosciences).

### END-seq

Detailed protocol for END-seq has been described elsewhere (Canela *et al*, 2016; Wong *et al*, 2021). Briefly, KM12 or HCT116 cells (10 million) along with the spike-in control (20%) were carefully resuspended in cell suspension buffer (BioRad). The resultant homogeneous single-cell suspensions were embedded in agarose and transferred into plug molds (Bio-Rad CHEF Mammalian Genomic DNA plug kit). Plugs were allowed to solidify at 4°C and were then incubated with Proteinase K (QIAGEN) (1 h at 50°C and then overnight at 37°C). The de-proteinized plugs were sequentially washed in Wash Buffer (10 mM Tris pH 8.0, 50 mM EDTA) and TE buffer (10 mM Tris pH 8.0, 1 mM EDTA), treated with RNaseA (QIAGEN), washed again in Wash Buffer. Blunting, A-tailing, and ligation of biotinylated hairpin adaptor 1 (ENDseq-adaptor-1, 5′-Phos-GATCGGAAGAGCGTCGTGTAGGGAAAGAGTGUU[Biotin-dT]U[Biotin-dT]UUACACTCTTT CCCTACACGACGCTCTTCCGATC*T-3′ [*phosphorothioate bond]) were performed in the plug. Thereafter, DNA was recovered by first melting the agarose plugs and then digesting the agarose with GELase (Epicenter), as per manufacturer’s recommendations. The recovered DNA was then sheared to a length between 150 and 200 bp by sonication (Covaris) followed by capture of biotinylated DNA fragments using streptavidin beads (MyOne C1, Invitrogen). The second end of the captured DNA were repaired using an enzymatic cocktail (15 U T4 DNA polymerase, 5 U Klenow fragment, 15 U T4 polynucleotide kinase), A-tailed with Klenow exo(-) (15U) and ligated to hairpin adaptor 2 (ENDseq-adaptor-2, 5′-Phos-GATCGGAAGAGCACACGTCUUUUUUUUAGACGTGTGCTCTTCCGATC*T-3′ [*phosphorothioate bond]) (Quick ligation kit, NEB). To prepare libraries for sequencing, the hairpins on both ENDseq adapters were digested with USER enzyme (NEB) and PCR amplified for 16 cycles using TruSeq index adapters. All libraries were quantified using Picogreen or qPCR. Sequencing was performed on the Illumina Nextseq500 (75 bp single-end reads). Alignment of reads and peak calling were performed as previously described (van Wietmarschen *et al*., 2020).

### Immunofluorescence

Cells were seeded on 12 mm round glass coverslips overnight and treated with DMSO or 0.5 μM dTAG-13 the following day for 24 h. Cells were additionally incubated with 10 μM EdU) during the last 30 min. Depending on the application, cells may be fixed immediately with 4% paraformaldehyde (10 min) or first pre-extracted (20 mM HEPES, 50 mM NaCl, 3 mM MgCl_2_, 0.3 M sucrose, 0.2% Triton X-100) on ice for 5 min to remove soluble nuclear proteins prior to fixation. Samples were then permeabilized (0.5% Triton X-100, 5 min) and blocked in 2% BSA/PBS. Incubation of primary antibodies recognizing RAD51 (1:250, Millipore), Cyclin A (1:200, Santa Cruz) and/or RPA2 (1:2,000, Cell Signaling) was followed by appropriate fluorochrome-conjugated secondary antibodies (Invitrogen). Next, click-IT chemistry was performed as per manufacturer’s instructions (Thermo Fisher Scientific) and DNA was counterstained with DAPI. Wide-field fluorescence images were captured at 40x magnification on a Lionheart LX automated microscope (BioTek Instruments). Quantification of nuclear foci was performed using the Gen5 spot analysis software (BioTek Instruments). Confocal z-stacks were acquired using a Nikon SoRa spinning disk microscope equipped with a 60x Apo TIRF oil immersion objective lens (N.A. 1.49) and Photometrics BSI sCMOS camera. Z-stacks were collected with 0.2 μm step size and 0.110 μm x-y pixel size. A denoise.ai algorithm was applied to the images in the dataset using the Nikon Elements (v.5.41) image analysis software. The denoised images were imported into Imaris image analysis software (Oxford Instruments) and the ImarisCell module was used to segment the individual nuclei and measure both the number of RAD51 foci per nucleus and the volume of each focus.

### Flow cytometry

Vehicle– and dTAG-13 treated cells were incubated with 10 μM EdU for 30 min at 37 °C, fixed with 1% formaldehyde for 10 min and stained using the Click-IT EdU Alexa Fluor 647 Flow Cytometry Assay Kit (Thermo Fisher Scientific) as per manufacturer’s instructions. DNA content was assessed by incorporation of DAPI (1 μg/ml). Where indicated, cells were first pulse-labeled with 5 μM CFSE (10 min at 37 °C) and treated or not with 0.5 μM dTAG-13 the following day. At 24-72 h post treatment, live cells were collected for FACS. Data was acquired on a Cytoflex tabletop flow cytometer (Beckman Coulter) and analyzed using FlowJo (BD Biosciences).

### Preparation and analysis of metaphase spreads

Cells were treated with DMSO or 0.5 μM dTAG-13 for 18 h and subsequently arrested at mitosis with 0.1 mg/ml colcemid (Roche) for 6 h (total 24 h). Metaphase chromosome spreads were prepared as previously described (Zong *et al*, 2019; Zong *et al*., 2015). First, cells were induced to swell in a prewarmed hypotonic solution containing 0.75 M KCl (Sigma) for 20 min at 37°C. Next, suspensions of single cells were fixed with a solution mixture containing methanol and glacial acetic acid at a 3:1 ratio. Thereafter, cells were further washed extensively with the same fixative solution and dropped onto slides in a humidified chamber (Thermotron Industries). To visualize metaphase chromosomes by fluorescence *in situ* hybridization (FISH), samples immobilized on slides were sequentially treated with pepsin (5-10 μg/ml in 0.01 N HCl, 5 min at 37°C), washed, dehydrated with ethanol, briefly heat denatured (80°C, 1 min 15 s) in a slide moat (Boekel Scientific) and incubated with a commercially available Cy3-labeled (CCCTAA)3 peptide nucleic acid probe (PNA Bio) recognizing mammalian telomere sequences. After extensive washes, DNA was counterstained with DAPI. Images were acquired at 63x magnification using the Metafer automated scanning and imaging platform (MetaSystems). Fifty individual metaphases were scored for the presence of chromosomal aberrations.

### CRISPR–Cas9 screen

CRISPR–Cas9 screen was performed using the whole genome human Brunello CRISPR knockout pooled library (Addgene #73178) (PMID: 26780180). RKO FKBP-WRN (GFP clone #17) stably expressing Cas9 were transduced at a multiplicity of infection (MOI) of 0.2 and 200-fold coverage of the library. The following day, cells were selected with puromycin (0.5 μg/mL) for three days and further expanded for four days. Aliquots of day zero cells were collected, and the remaining cells were split into triplicates for drug treatment (DMSO vs 0.5 μM dTAG-13), maintaining 200-fold coverage for a further six days. Surviving cells from each condition were collected, and genomic DNA was isolated (Blood & Cell Culture DNA Midi Kit, Qiagen). DNA was PCR amplified with Illumina-compatible primers (P7 and P5 mix), gel purified and processed for Illumina sequencing. Genes enriched or depleted in the dTAG-13 treated samples were determined with the MAGeCK software package version 0.5.9.5.

### Growth competition assay

Sequences for sgRNAs of interest were cloned into LRG2.1-GFP (Addgene #108098) (Tarumoto *et al*, 2018) as described (Girish & Sheltzer, 2020) and Sanger sequenced to confirm correct insertion. RKO FKBP-WRN (GFP clone #17, Puro clone #16) and KM12 FKBP-WRN (Puro clone #5) stably expressing Cas9 were seeded and transduced with LRG2.1-GFP containing the sgRNA of interest (MOI approximately 0.5). After three days, cells were split into control and dTAG-13 (0.5 μM) and an aliquot was run on flow cytometry to establish GFP expression at baseline. On the indicated days, an aliquot of cells was collected for flow cytometry analysis and the remaining cells were split and maintained in dTAG-13 as needed. To verify knockdown of target genes, cells treated or not with 0.5 μM dTAG-13 for 24 h before collection. RNA was extracted (RNeasy Mini Kit, Qiagen) and 1 μg was used to make cDNA (SuperScript VILO cDNA Synthesis Kit, Invitrogen). Quantitative RT-PCR was performed with iTAQ Syber Green using 1/40th of the cDNA reaction per sample and the appropriate primers. Western blotting was performed as described above.

### Statistical analyses

Unless otherwise indicated, statistical significance was evaluated with unpaired Student’s t-test or Mann-Whitney nonparametric test. The resultant P values (represented by asterisks) are indicated in the figure panels and were calculated using GraphPad Prism (ns = p > 0.05; ∗ = p < 0.05; ∗∗ = p < 0.01; ∗∗∗ = p < 0.001; ∗∗∗ = p < 0.0001). The total number of replicates, mean and error bars are explained in the figure legends.

## Acknowledgment

We thank Drs. Eros Lazzerini Denchi, Sergio Ruiz, Michael Lichten, Sam John and members of the Laboratory of Genome Integrity for discussions and advice; Drs. Ferenc Livak, Subhadra Banerjee and Shafiuddin Siddiqui at the CCR/LGI Flow Cytometry Core for cell sorting; and the NCI/CCR Genomics Core for help with sequencing. This work used the computational resources of the NIH HPC Biowulf cluster.

## Figure legends

**Appendix Fig S1. Verification of FKBP-WRN knock-in at the endogenous WRN locus.** (A) Schematics depicting the locations of the two sgRNAs (in blue and red) used to target the WRN gene. The translation initiation site of WRN is indicated by the green underlined ATG codon. (B) Confirmation of successful FKBP-WRN knock-in. Genomic DNA was extracted from candidate clones and amplified by PCR prior to Sanger sequencing. Note that both homozygous and heterozygous FKBP-WRN knock-in clones were obtained. In the case of heterozygous knock-in, the remaining allele of WRN was found to be inactivated by small indels that either excised the initiation ATG or produced a STOP codon slightly downstream of the ATG.

**Appendix Fig S2. Short-term DNA damage responses in MSI-H cancer cells following acute WRN degradation.** (A) RKO and KM12 clones expressing FKBP-WRN were treated or not with 0.5 μM dTAG-13 for the indicated amount of time. Immunoblotting showed the levels of pCHK2, CHK1 and RPA2 as a function of time. One of two independent experiments is shown. (B) RKO c16 was treated or not with 0.5 μM dTAG-13 for 4 or 24 h. Where indicated, 3 mM HU and/or 1 μM AZ-20 was added to the growth media for 2 h. Immunoblotting showed that ATR autophosphorylation, ATR-mediated CHK1 (S345) and RPA (S33) phosphorylation and CHK1 autophosphorylation (S296) were induced by HU regardless of the WRN status. A representative experiment is shown. (C) RKO c16 was treated or not with 0.5 μM dTAG-13 for 24 h or γ-irradiated with 5 Gy and allowed to recover for 4 h. Immunofluorescence staining showed enlarged RAD51 foci in dTAG-treated G2 cells, as compared to RAD51 foci found in S-phase cells. By contrast, irradiation induced RAD51 foci of comparable sizes in S and G2. One of two independent experiments is shown.

**Appendix Fig S3. Long-term DNA damage responses in MSI-H cancer cells following acute WRN degradation.** (A) RKO, KM12 and HCT116 clones expressing FKBP-WRN were pre-labeled with 5 μM CFSE and then treated or not with 0.5 μM dTAG-13. FACS analyses of live cells at the indicated post-treatment times showed that dTAG-treated MSI-H cancer cells could still divide, albeit at a reduced rate compared to vehicle-treated counterparts. Average of three (RKO, HCT116) or two (KM12) independent experiments is shown. (B) RKO clones expressing FKBP-WRN were treated or not with 0.5 μM dTAG-13 or with 1 μg/mL doxycycline for the indicated amounts of time. Immunoblotting showed progressive induction of CHK2 phosphorylation, but not that of CHK1. Total CHK1 protein level remained unchanged over time. One of two independent experiments is shown. (C) KM12 and HCT116 clones expressing FKBP-WRN were treated or not with 0.5 μM dTAG-13 for the indicated amounts of time. Immunoblotting showed progressive induction of pCHK2 over time. Phosphorylation of CHK1 was also induced but did not seem to increase over time.

**Appendix Fig S4. Lack of evidence for *bona fide* WRN resistance genes.** (A) RKO c17/Cas9 cells were transduced with individual sgRNAs targeting candidate WRN resistance genes identified from the CRISPR screen or a control sgRNA (ROSA) and subjected to growth competition assay (see Fig. 5D). Aliquots of vehicle– and dTAG-treated cells were taken after 24 h and the knockdown efficiency of target genes were determined by qRT-PCR. (B) Similar to (A), except that aliquots of vehicle– and dTAG-treated cells transduced with individual sgRNAs targeting candidate WRN resistance genes were processed for immunoblotting. Note that cells depleted for ILF3 were no longer able to efficiently degrade FKBP-WRN in response to dTAG-13. (C) A different clone of RKO (c16/Cas9) was transduced with individual sgRNAs targeting a subset of candidate WRN resistance genes and subjected to growth competition assays. (D) A clone of KM12 (c5/Cas9) was transduced with individual sgRNAs targeting a subset of candidate WRN resistance genes and subjected to growth competition assays. Note that only depletion of CRBN reproducibly rescued lethality in both dTAG-treated RKO and KM12 clones. A representative experiment is shown for each figure panel.

**Appendix Fig S5. Dose-dependent cytotoxicity imparted by dTAG-13 and AZ-20 in human MSI-H cancer cells.** (A-C) RKO (A), KM12 (B) and HCT116 (C) clones expressing FKBP-WRN were treated or not with indicated doses of dTAG-13. After 24 h, cells were collected for immunoblotting (upper panels). A representative blot is shown. Cell viability was measured after 6 days (lower panels). Average (+/-S.D.) of six independent experiments is shown. (D) A larger panel of human MSI-H and MSS colorectal cancer cell lines were treated or not with indicated doses of AZ-20. Cell viability was measured after 6 days. Average (+/-S.D.) of three (RKO, KM12, LS180, SW48, NCIH508, SW620, SW837) to five (HCT116) independent experiments is shown. (E) MSI-H (RKO, KM12, HCT116 and MSS (NCIH508, SW837) cancer cells were treated or not with 0.5 μM AZ-20 for 24 h (+ colcemid for the last 6 h). Analysis of metaphase spreads showed that ATR inhibition induced high levels of chromosomal aberrations in MSI-H but not MSS cells. Average (+/-S.D.) of three independent experiments. (F) The impact of WRN and ATR depletion on the fitness of human MSI-H vs MSS cancer cells (colorectal + endometrial). Publicly available data was downloaded from DepMap and re-plotted. (G) The cytotoxicity profiles of three ATR inhibitors in human MSI-H vs MSS cancer cells (colorectal + endometrial). A.U.C. values were downloaded from Genomics of Drug Sensitivity in Cancer (GDSC) or DepMap (PRISM repurposing screens) and re-plotted.

**Appendix Fig S6. ATR co-inhibition enhances the cytotoxicity of low dose dTAG-13 treatment.** (A) RKO clones expressing FKBP-WRN were treated or not with the indicated doses of dTAG-13, AZ-20 or their combination. Average (+/-S.D.) of four independent experiments is shown. Cell viability was measured after 6 days. (B) Similar to (A) but in HCT116 and KM12 clones. Average (+/-S.D.) of four independent experiments is shown. (C) HCT116 (left panel) and KM12 (right panel) clones expressing FKBP-WRN were treated or not with the indicated doses of dTAG-13, AZ-20 or their combination for 24 h. END-seq showed that the combination regimen resulted in elevated levels of genomic breakage at expanded TA repeat regions, as compared to each single agent treatment.

**Appendix Movie S1. Examples of dTAG-induced cell death events.** RKO reporter cells were treated 0.5 μM dTAG-13 for and tracked by time-lapse microscopy. The events shown in this movie occurred between frames 270-310, which corresponded to 54-62 h after drug treatment. Note that during death, the nucleus could violently disintegrate or remain relatively intact.

## References

1. Adikusuma F, Pfitzner C, Thomas PQ (2017) Versatile single-step-assembly CRISPR/Cas9 vectors for dual gRNA expression. PLoS One 12: e0187236

2. Ammazzalorso F, Pirzio LM, Bignami M, Franchitto A, Pichierri P (2010) ATR and ATM differently regulate WRN to prevent DSBs at stalled replication forks and promote replication fork recovery. EMBO J 29: 3156–3169

3. Behan FM, Iorio F, Picco G, Goncalves E, Beaver CM, Migliardi G, Santos R, Rao Y, Sassi F, Pinnelli M et al (2019) Prioritization of cancer therapeutic targets using CRISPR-Cas9 screens. Nature 568: 511–516

4. Bryant HE, Schultz N, Thomas HD, Parker KM, Flower D, Lopez E, Kyle S, Meuth M, Curtin NJ, Helleday T (2005) Specific killing of BRCA2-deficient tumours with inhibitors of poly(ADP-ribose) polymerase. Nature 434: 913–917

5. Canela A, Sridharan S, Sciascia N, Tubbs A, Meltzer P, Sleckman BP, Nussenzweig A (2016) DNA Breaks and End Resection Measured Genome-wide by End Sequencing. Mol Cell 63: 898–911

6. Cappell SD, Chung M, Jaimovich A, Spencer SL, Meyer T (2016) Irreversible APC(Cdh1) Inactivation Underlies the Point of No Return for Cell-Cycle Entry. Cell 166: 167–180

7. Chan EM, Shibue T, McFarland JM, Gaeta B, Ghandi M, Dumont N, Gonzalez A, McPartlan JS, Li T, Zhang Y et al (2019) WRN helicase is a synthetic lethal target in microsatellite unstable cancers. Nature 568: 551–556

8. Chari R, Yeo NC, Chavez A, Church GM (2017) sgRNA Scorer 2.0: A Species-Independent Model To Predict CRISPR/Cas9 Activity. ACS Synth Biol 6: 902–904

9. Dimitrova N, Chen YC, Spector DL, de Lange T (2008) 53BP1 promotes non-homologous end joining of telomeres by increasing chromatin mobility. Nature 456: 524–528

10. Farmer H, McCabe N, Lord CJ, Tutt AN, Johnson DA, Richardson TB, Santarosa M, Dillon KJ, Hickson I, Knights C et al (2005) Targeting the DNA repair defect in BRCA mutant cells as a therapeutic strategy. Nature 434: 917–921

11. Girish V, Sheltzer JM (2020) A CRISPR Competition Assay to Identify Cancer Genetic Dependencies. Bio Protoc 10: e3682

12. Giunta S, Belotserkovskaya R, Jackson SP (2010) DNA damage signaling in response to double-strand breaks during mitosis. J Cell Biol 190: 197–207

13. Grant GD, Kedziora KM, Limas JC, Cook JG, Purvis JE (2018) Accurate delineation of cell cycle phase transitions in living cells with PIP-FUCCI. Cell Cycle 17: 2496–2516

14. Hao S, Tong J, Jha A, Risnik D, Lizardo D, Lu X, Goel A, Opresko PL, Yu J, Zhang L (2022) Synthetical lethality of Werner helicase and mismatch repair deficiency is mediated by p53 and PUMA in colon cancer. Proc Natl Acad Sci U S A 119: e2211775119

15. Jin Z, Sinicrope FA (2022) Mismatch Repair-Deficient Colorectal Cancer: Building on Checkpoint Blockade. J Clin Oncol 40: 2735–2750

16. Kategaya L, Perumal SK, Hager JH, Belmont LD (2019) Werner Syndrome Helicase Is Required for the Survival of Cancer Cells with Microsatellite Instability. iScience 13: 488–497

17. Li W, Xu H, Xiao T, Cong L, Love MI, Zhang F, Irizarry RA, Liu JS, Brown M, Liu XS (2014) MAGeCK enables robust identification of essential genes from genome-scale CRISPR/Cas9 knockout screens. Genome Biol 15: 554

18. Lieb S, Blaha-Ostermann S, Kamper E, Rippka J, Schwarz C, Ehrenhofer-Wolfer K, Schlattl A, Wernitznig A, Lipp JJ, Nagasaka K et al (2019) Werner syndrome helicase is a selective vulnerability of microsatellite instability-high tumor cells. Elife 8

19. Morales-Juarez DA, Jackson SP (2022) Clinical prospects of WRN inhibition as a treatment for MSI tumours. NPJ Precis Oncol 6: 85

20. Nabet B, Roberts JM, Buckley DL, Paulk J, Dastjerdi S, Yang A, Leggett AL, Erb MA, Lawlor MA, Souza A et al (2018) The dTAG system for immediate and target-specific protein degradation. Nat Chem Biol 14: 431–441

21. Nelson G, Buhmann M, von Zglinicki T (2009) DNA damage foci in mitosis are devoid of 53BP1. Cell Cycle 8: 3379–3383

22. Olave MC, Graham RP (2022) Mismatch repair deficiency: The what, how and why it is important. Genes Chromosomes Cancer 61: 314–321

23. Pettitt SJ, Krastev DB, Brandsma I, Drean A, Song F, Aleksandrov R, Harrell MI, Menon M, Brough R, Campbell J et al (2018) Genome-wide and high-density CRISPR-Cas9 screens identify point mutations in PARP1 causing PARP inhibitor resistance. Nat Commun 9: 1849

24. Picco G, Cattaneo CM, van Vliet EJ, Crisafulli G, Rospo G, Consonni S, Vieira SF, Rodriguez IS, Cancelliere C, Banerjee R et al (2021) Werner Helicase Is a Synthetic-Lethal Vulnerability in Mismatch Repair-Deficient Colorectal Cancer Refractory to Targeted Therapies, Chemotherapy, and Immunotherapy. Cancer Discov 11: 1923–1937

25. Pichierri P, Rosselli F, Franchitto A (2003) Werner’s syndrome protein is phosphorylated in an ATR/ATM-dependent manner following replication arrest and DNA damage induced during the S phase of the cell cycle. Oncogene 22: 1491–1500

26. Schindelin J, Arganda-Carreras I, Frise E, Kaynig V, Longair M, Pietzsch T, Preibisch S, Rueden C, Saalfeld S, Schmid B et al (2012) Fiji: an open-source platform for biological-image analysis. Nat Methods 9: 676–682

27. Spencer SL, Cappell SD, Tsai FC, Overton KW, Wang CL, Meyer T (2013) The proliferation-quiescence decision is controlled by a bifurcation in CDK2 activity at mitotic exit. Cell 155: 369–383

28. Taieb J, Svrcek M, Cohen R, Basile D, Tougeron D, Phelip JM (2022) Deficient mismatch repair/microsatellite unstable colorectal cancer: Diagnosis, prognosis and treatment. Eur J Cancer 175: 136–157

29. Tarumoto Y, Lu B, Somerville TDD, Huang YH, Milazzo JP, Wu XS, Klingbeil O, El Demerdash O, Shi J, Vakoc CR (2018) LKB1, Salt-Inducible Kinases, and MEF2C Are Linked Dependencies in Acute Myeloid Leukemia. Mol Cell 69: 1017–1027 e1016

30. van Wietmarschen N, Nathan WJ, Nussenzweig A (2021) The WRN helicase: resolving a new target in microsatellite unstable cancers. Curr Opin Genet Dev 71: 34–38

31. van Wietmarschen N, Sridharan S, Nathan WJ, Tubbs A, Chan EM, Callen E, Wu W, Belinky F, Tripathi V, Wong N et al (2020) Repeat expansions confer WRN dependence in microsatellite-unstable cancers. Nature 586: 292–298

32. Waterman DP, Haber JE, Smolka MB (2020) Checkpoint Responses to DNA Double-Strand Breaks. Annu Rev Biochem 89: 103–133

33. Wong N, John S, Nussenzweig A, Canela A (2021) END-seq: An Unbiased, High-Resolution, and Genome-Wide Approach to Map DNA Double-Strand Breaks and Resection in Human Cells. Methods Mol Biol 2153: 9–31

34. Zong D, Adam S, Wang Y, Sasanuma H, Callen E, Murga M, Day A, Kruhlak MJ, Wong N, Munro M et al (2019) BRCA1 Haploinsufficiency Is Masked by RNF168-Mediated Chromatin Ubiquitylation. Mol Cell 73: 1267–1281 e1267

35. Zong D, Callen E, Pegoraro G, Lukas C, Lukas J, Nussenzweig A (2015) Ectopic expression of RNF168 and 53BP1 increases mutagenic but not physiological non-homologous end joining. Nucleic Acids Res 43: 4950–4961

