## Supplementary figures and images for "Comprehensive mapping of cell fates in microsatellite unstable cancer cells support dual targeting of WRN and ATR"

### Supplementary figures 1-6

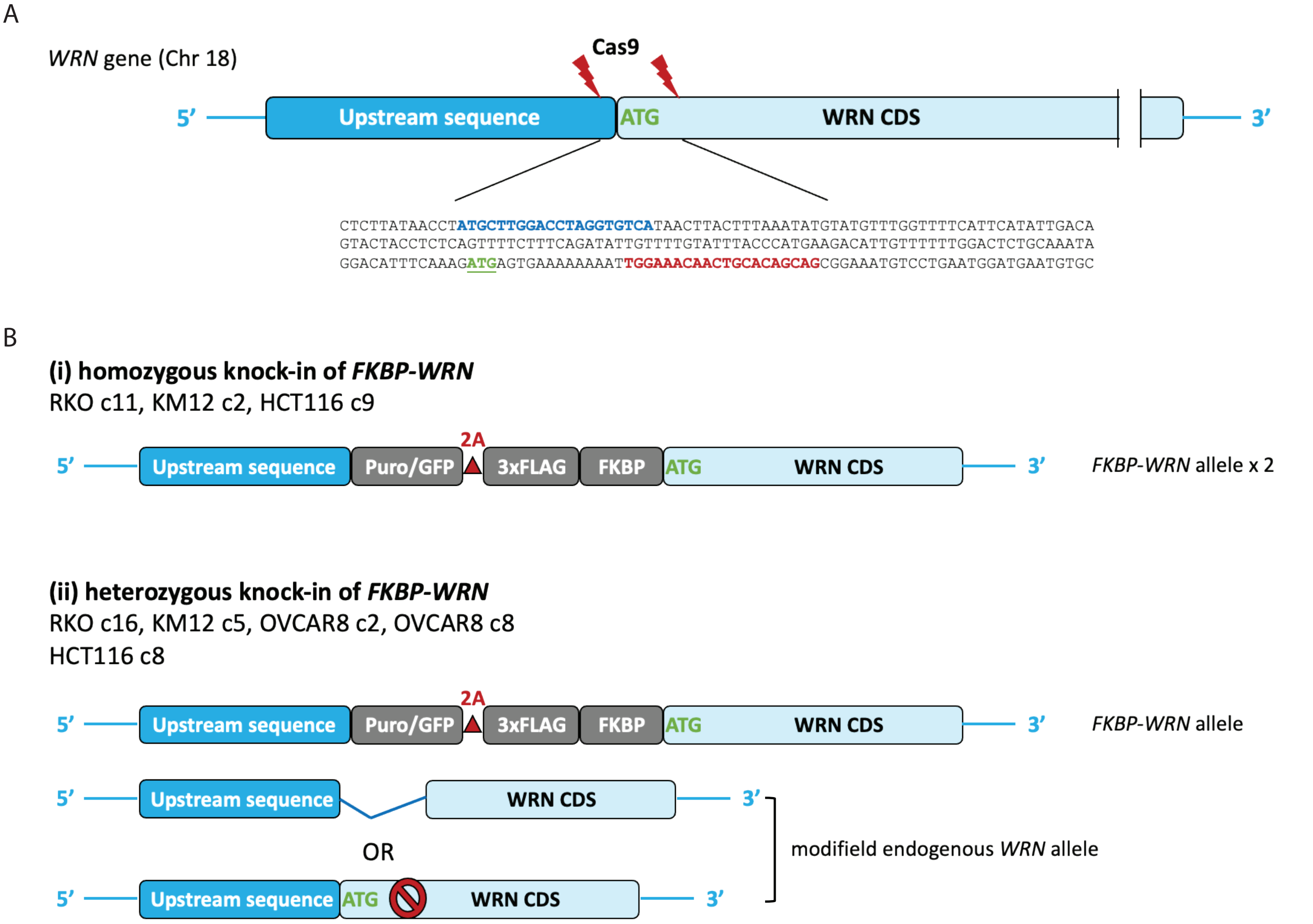

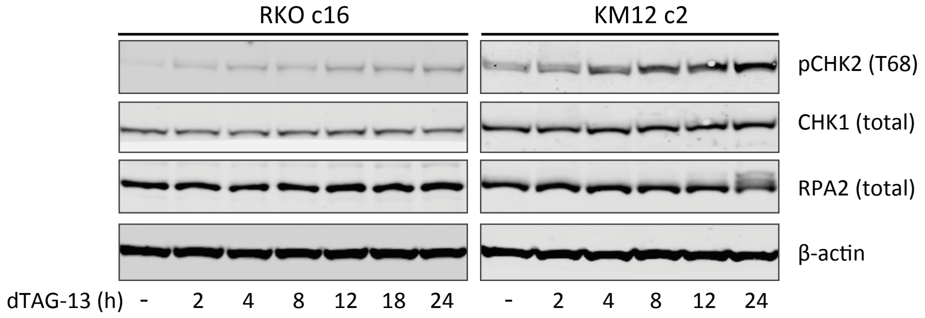

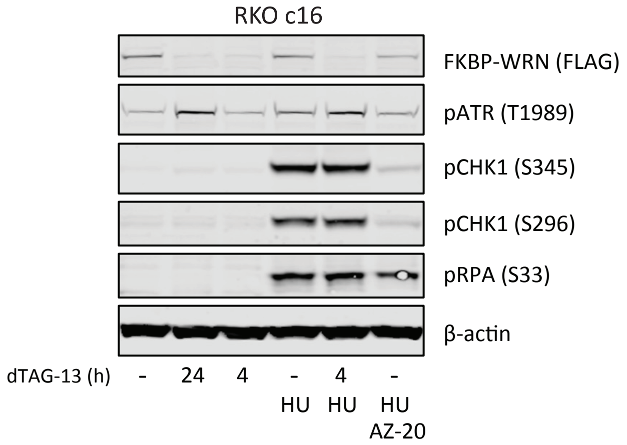

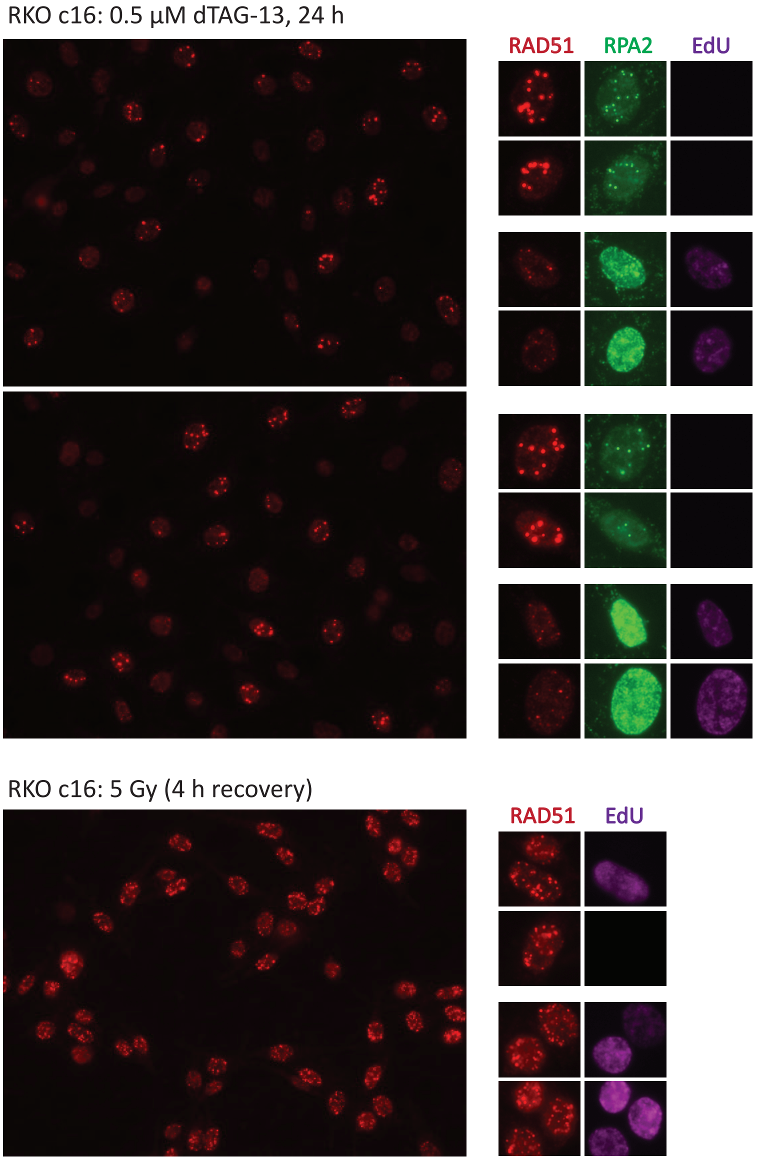

Appendix Fig S3

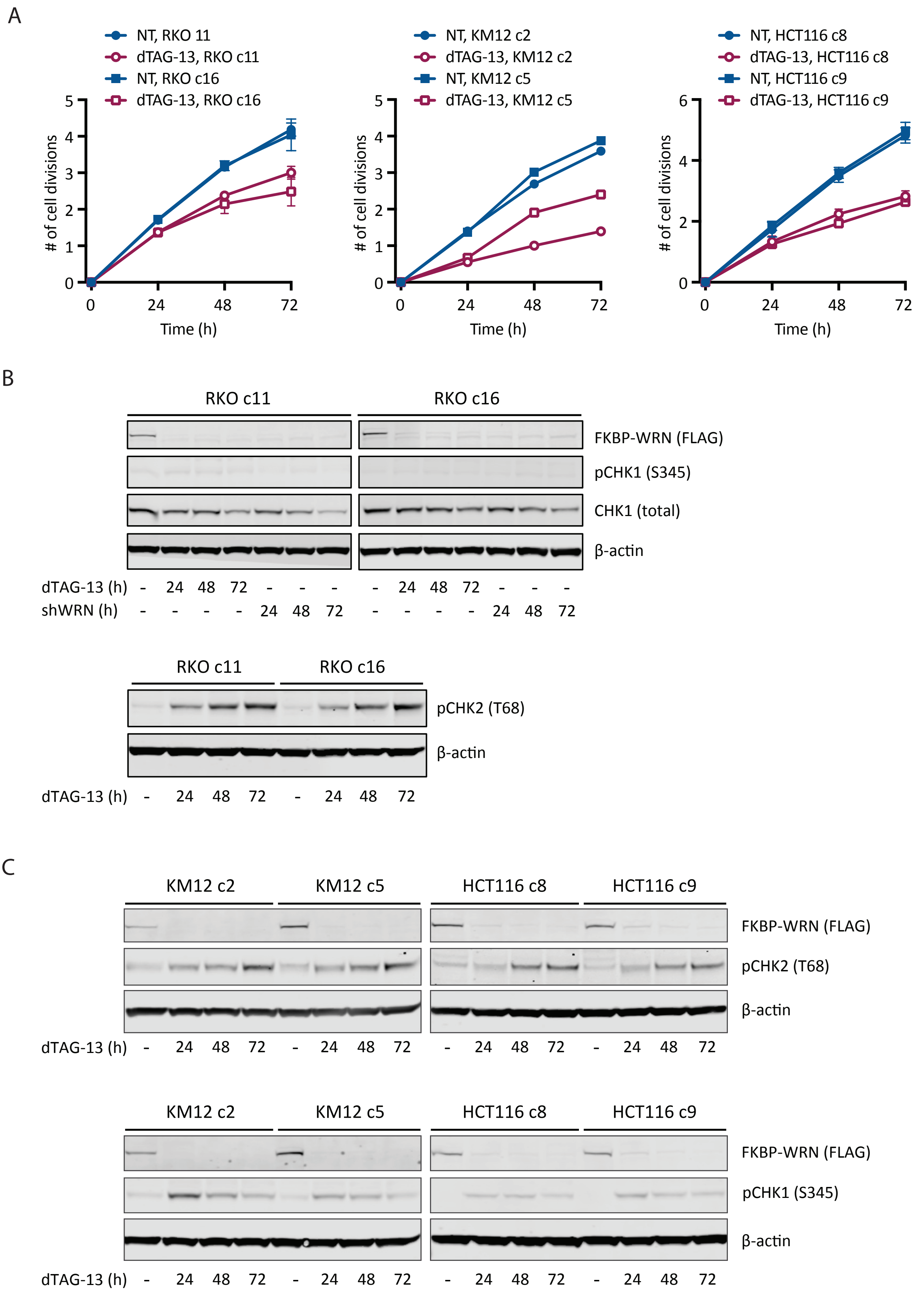

Appendix Fig S4

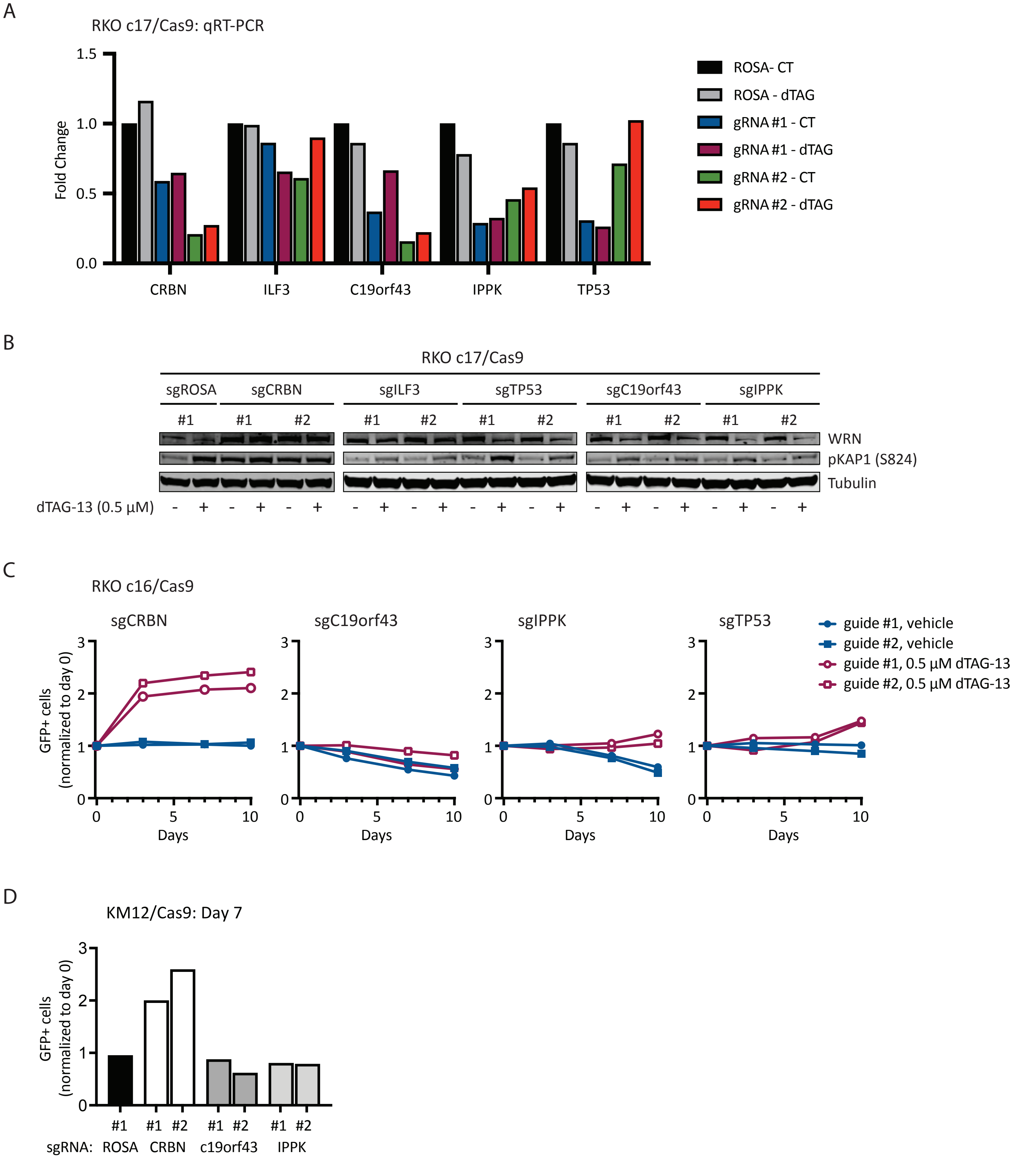

Appendix Fig S5

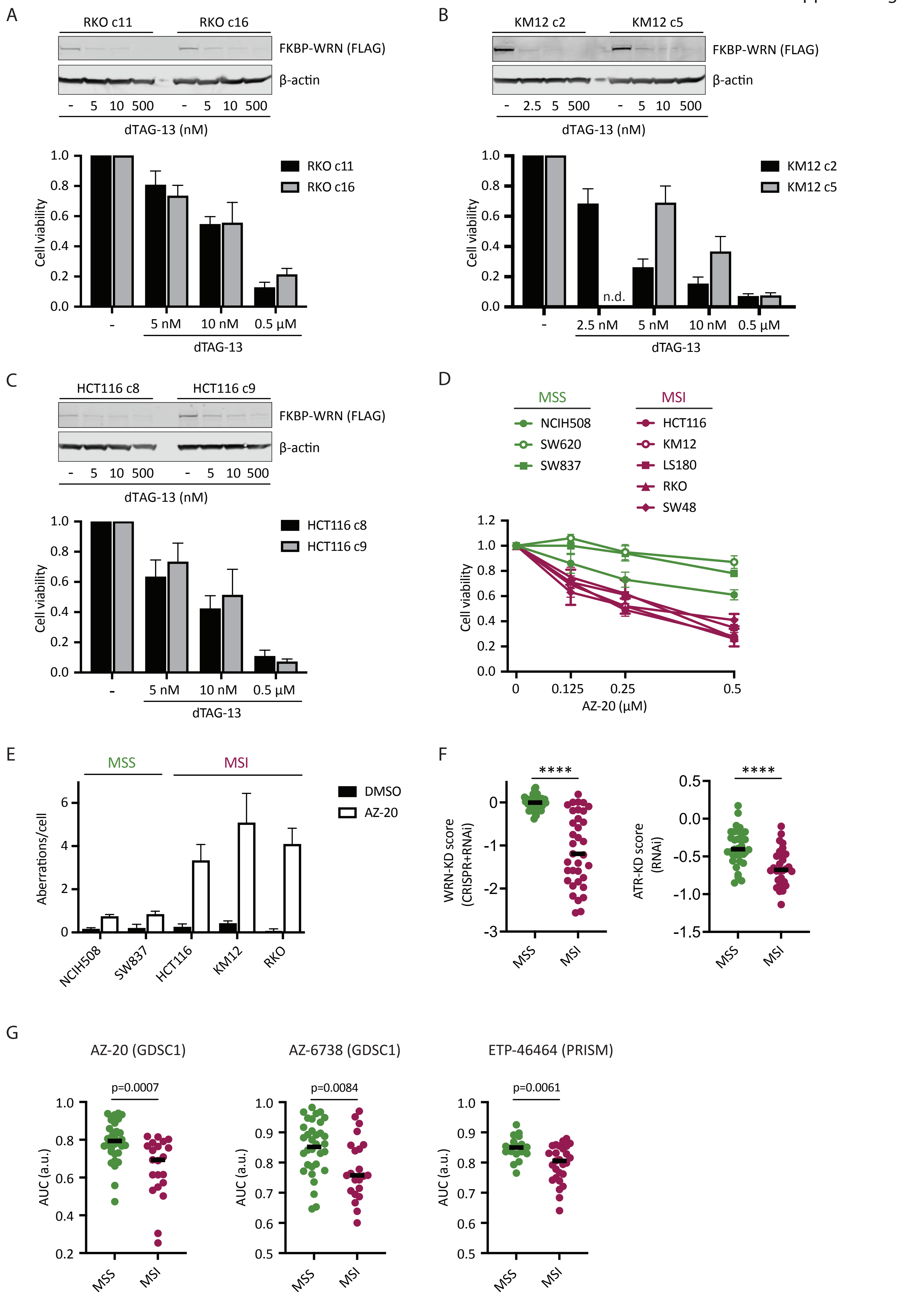

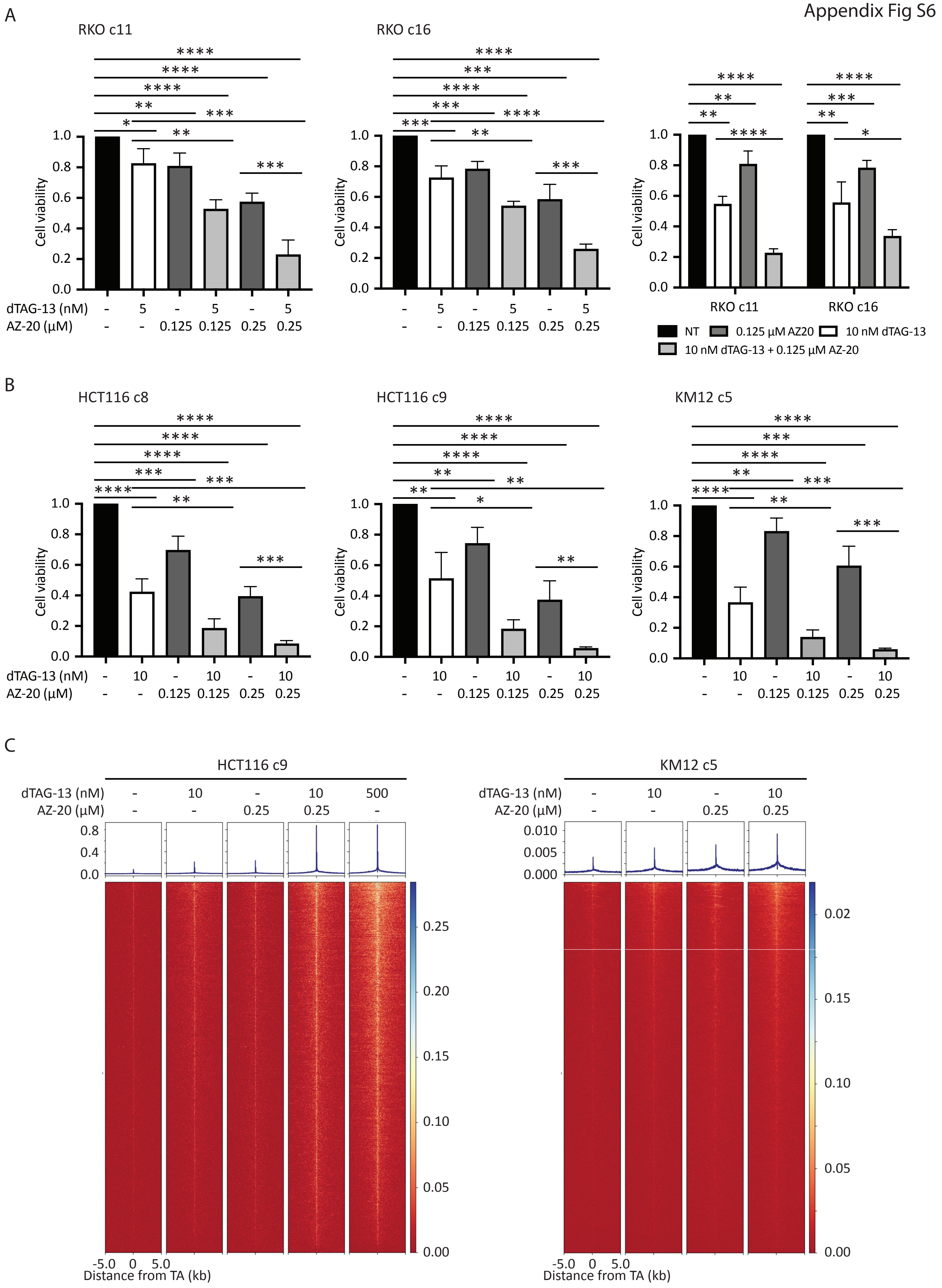
